# Intussusceptive angiogenesis-on-a-chip: Evidence for transluminal vascular bridging by endothelial delamination

**DOI:** 10.1101/2024.12.19.627594

**Authors:** Sabrina C.R. Staples, Hao Yin, Frances S.K. Sutherland, Emma Prescott, Dylan Tinney, Douglas W. Hamilton, Daniel Goldman, Tamie L. Poepping, Christopher G. Ellis, J. Geoffrey Pickering

## Abstract

Intussusceptive angiogenesis is an increasingly recognized vessel duplication process that generates and reshapes microvascular beds. However, the mechanism by which a vessel splits into two is poorly understood. Particularly vexing is formation of the hallmark transluminal endothelial cell bridge. How an endothelial cell comes to cross a flowing lumen rather than line it is enigmatic. To elucidate this, we used a microvessel-on-a-chip strategy, creating a micro-conduit coherently lined with flow-sensitive endothelial cells but in which transluminal bridges also formed. Bridge morphologies ranged from filamentous strand to multicellular columns with a central core. These bridge architectures were found to recapitulate those in microvessels in embryos, tumours, diseased organs, and the dermis of patients with limb-threatening ischemia. Time-lapse, multi-plane, 3D microscopy of the micro-physiologic conduit revealed that bridges arose from endothelial cells oriented orthogonal to flow that partially released from the wall while retaining attachments at the ends. This delamination process was blocked by hyperactivation of Rho and augmented by interventions that weaken cell-substrate interactions, including inhibiting non-muscle myosin II and blocking α5ß1 integrin but, interestingly, not αvß3 integrin. Thus, endothelial cells can leave their monolayer and transect a flowing lumen through controlled delamination. This previously unrecognized lumen entry program could explain the launch of intussusceptive angiogenesis and opens a framework for intervening.

**Significance Statement:** Rapid generation of small blood vessels is vital for embryonic development and many diseases. An efficient means of creating a new microvessel is for an existing vessel to split into two, a process recognized for over 35 years. However, the cellular events underlying vessel splitting remain largely a mystery. The challenge is how to look inside a microvessel and capture transient events. We address this challenge using a microvessel-on-a-chip strategy. We discovered that select endothelial cells lining the wall can partially lift to transect the lumen. This resolves the paradox of an adhesion-dependent cell reaching across a pressurized lumen. It also reframes microvascular therapy considerations, including for diabetics with leg ulcers, a condition we show has hallmarks of microvessel splitting.

## Introduction

Angiogenesis is fundamental to embryonic development, tissue regeneration, and many diseases (1–3). The best studied mode of angiogenesis is sprouting angiogenesis. However, intussusceptive angiogenesis (IA), a process characterized by the longitudinal division of a vessel into two daughter vessels, is increasingly recognized.

There are many contexts for IA including the developing lung, myocardium, liver, and intestines (4–6). IA is also important in adults. For example, in adult skeletal muscle IA mediates the vascular network expansion response to increased blood flow (7) and is also the major mode of angiogenesis after ischemic injury (8). IA also proceeds in tumours (9–11) where its emergence may confound anti-angiogenesis strategies (12). Other disease contexts include colitis (13) and COVID-19 lung disease (14). Yet despite the growing recognition of IA, the mechanisms by which a vessel splits and duplicates are poorly understood (3).

A particularly enigmatic feature of IA is the endothelial transluminal bridge. This peculiar structure is comprised of one or more endothelial cells that span the lumen. Endothelial bridges are considered to be a precursor to the transluminal pillar, which has the added complexity of a tissue core which advances the lumen division process. Bridges and pillars are thus hallmarks of IA but they can also exist in pathological contexts without robust IA. For example, endothelial bridges that fail to progress to vessel splitting have been found to obstruct blood flow in multi-cavernous vascular malformations (15). Bridges and pillars have also been associated with microthrombi, as identified in the hearts of individuals succumbing to COVID-19 (14, 16, 17).

Understanding how transluminal bridges form has been challenging. Electron microscopy studies of lung capillaries have suggested that opposing endothelial cell surfaces can invaginate toward each other, make contact, and reorganize with adjacent stroma to form a pillar (5, 18). Capillary wall folding (19, 20) and incorporating an extraluminal vessel loop into a venous wall (20) are other processes described in embryonic vessels. However, IA has been abundantly seen in vessels larger than capillaries, including up to 100 µm in diameter, where opposing surface contact mechanisms are less likely (6, 8, 21, 22). Another proposal holds that the bridging proceeds via an inverse endothelial sprouting mechanism (13, 23). Although intriguing, the steps enabling such inverse sprouting are unknown and whether endothelial cells can generate the protrusive forces necessary to cross a pressurized, flowing lumen is questioned (24).

The limited understanding of endothelial cell bridge formation is not surprising given the difficulties in interrogating transient events inside a lumen. Corrosion casting, serial sectioning, and intravital microscopy can identify bridges and pillars (8, 25) but cannot directly inform on their dynamics. Live cell imaging holds promise but has been limited to the embryonic fish caudal vein plexus (20). Tracking intraluminal bridging in mammalian vessels remains a hurdle.

It thus is noteworthy that a major gap in studying IA is a relative lack of *in vitro* models. For sprouting angiogenesis culture models have been invaluable (26, 27) and they continue to advance the field (28–30). *In vitro* models of IA could be similarly powerful by enabling high resolution imaging and a platform for deciphering mechanisms, as well as offering cost efficiencies and opportunities for wider study among the research community. Because the processes of IA are fundamentally intralumenal, models with a closed lumen and fluid flow are needed (31). Recently, we made the first steps toward this goal using a microfluidic approach (8). However, the central question as to how one or more endothelial cells, which are adhesion-dependent cells, come to transect a flowing lumen rather than line it remains a mystery.

Herein, we investigate the dynamics of transluminal bridging using a microvessel-on-a-chip approach. We identify a diversity of endothelial bridge morphologies that recapitulate those found *in vivo*, including in patients with limb-threatening ischemia. Furthermore, we report a previously unrecognized endothelial cell delamination program, a controlled cell lifting process that underlies transluminal bridge formation.

## Results

### Generation of an endothelial microfluidic conduit capable of flow-mediated remodeling

To investigate endothelial cell dynamics in a vessel-on-a-chip environment, we generated a closed, small caliber (80 µm-diameter) 3D microchannel fully lined by endothelial cells.

Because of recognized challenges in maintaining healthy endothelial cells in a channel this small, we seeded endothelial cells onto a fibronectin-coated surface using a controlled, direct visualization cell delivery approach. Flow was then applied to the microchannel to yield wall shear stresses ranging from 0.04 to 20 dyn/cm^2^, recognizing that IA has been identified in diverse shear stress environments (8, 32).

A confluent monolayer of endothelial cells lining the entire length of the channel was found under all flow conditions (Fig. 1A). The endothelial cells formed VE-cadherin-rich cell-cell junctions and displayed abundant actin microfilament bundles (Fig. 1B). To assess the responsivity of the endothelial cells to fluid flow, polarity was assessed based on positioning of the Golgi apparatus. At 10 dyn/cm^2^ fluid shear stress, most endothelial cells (52±3%) polarized against the direction of flow (p<0.0001, Fig. 1C), consistent with *in vivo* data (33, 34). At 0.8 dyn/cm^2^, against-flow polarization still predominated (45±3%, p<0.0001 vs orthogonal, P<0.022 vs with-flow) although the distribution of polarization was wider (p=0.023, Fig. 1C). There was prominent alignment of actin microfilament bundles with flow under both flow conditions. This including actin bundles with a near-contiguous cell-to-cell appearance (Fig. 1B arrows).

**Figure 1.**
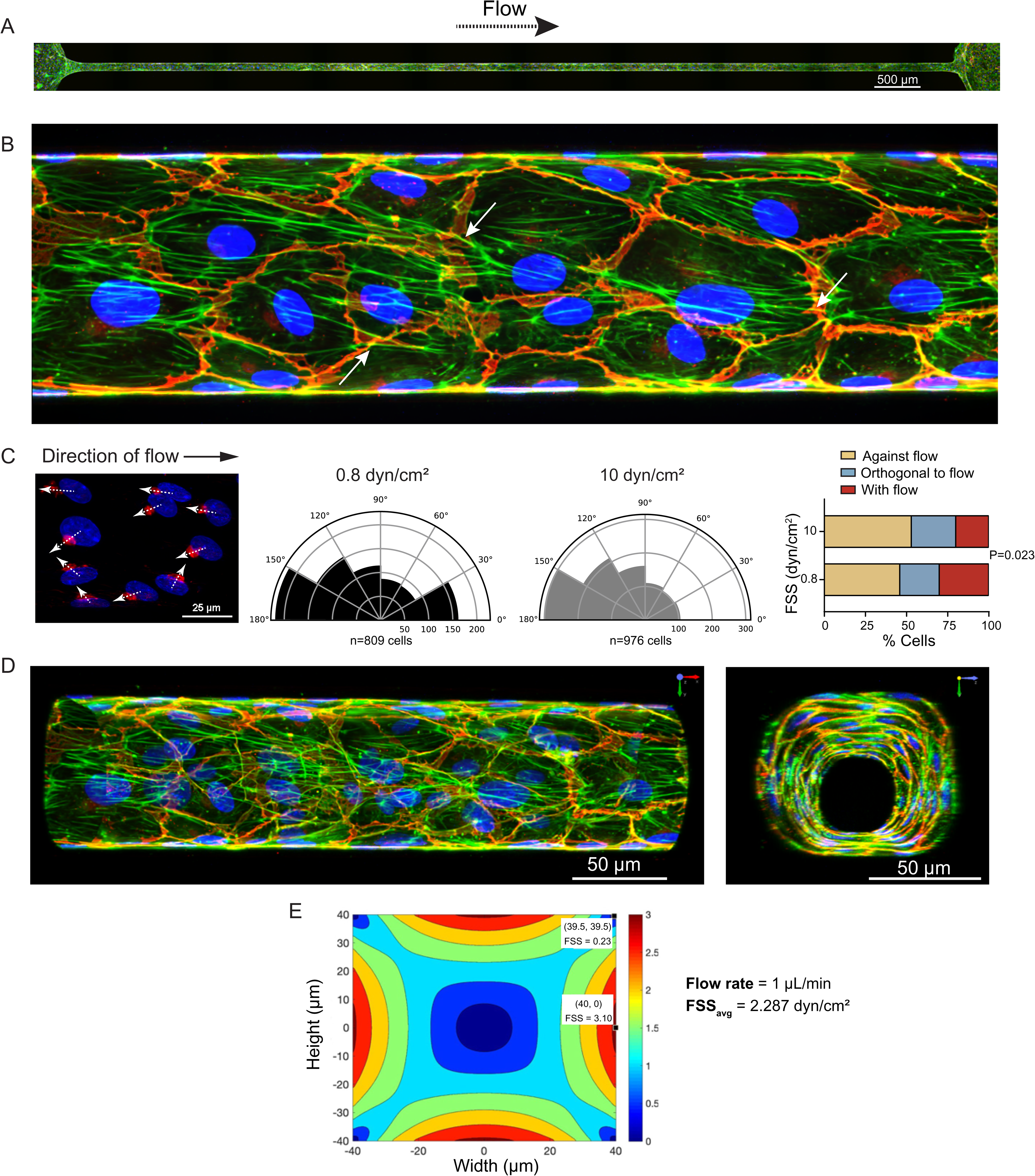
Endothelial cells form a confluent circumferential lining within PDMS channel. **A**. Confocal planar projection (50 z-slices, 2-µm step size) of 1 cm long HUVEC-lined microfluidic channel. Cells were incubated in static conditions for 16 h after seeding, thereafter subjected to flow at 2 dyn/cm^2^ for 72 hours. Cells are stained with Alexa Fluor-conjugated phalloidin to detect F-actin (green), DAPI for nuclear detection (blue), and immunostained for VE-cadherin (red). Scale bar, 500 µm. **B**. Higher magnification of a confocal planar projection (100 z-slices, 0.5 µm step size) of HUVECs residing on the bottom surface of a channel subjected to a shear stress of 2 dyn/cm^2^. There is a confluent monolayer with robust, predominantly linear, VE-cadherin junctions and actin stress fibres aligned in the direction of flow. **C**. *Left:* Fluorescent micrograph of endothelial cells immunostained for the Golgi apparatus (anti-GM130) and counterstained with DAPI. Arrows depict the cell polarity orientation, based on the Golgi-nucleus axis. *Middle:* Polar plots depicting the cell polarity relative to flow for endothelial cells subjected for 72 h to a shear stress of either 0.8 (809 cells) or 10 (976 cells) dyn/cm^2^. Cell number is depicted on the x-axis. *Right:* Graph depicting the distributions of cell polarity relative to flow. The distributions differ between flow conditions but, for both, polarization is biased against the flow (P=0.0001). **D.** 3D volume projections of an endothelialized microchannel in the x-y (left) and y-z (right) orientations. Scale bars correspond to dimensions of the front plane of the volume image. **E**. Fluid shear stress map derived from computational analysis of an 80 x 80 µm channel subjected to flow of 1 µL/min. The average shear stress was 2.3 dyn/cm^2^ but ranged from 3.10 dyn/cm^2^ at the mid-wall to as low as 0.23 dyn/cm^2^ at the corner. FSS, flow shear stress.

Remarkably, although the PDMS channels had a square cross section, 3D confocal reconstruction of the channels revealed inward remodeling of the endothelial conduit with rounding of microvessel corners that could yield a near-circular lumen cross section (Fig. 1D). To determine if this remodeling response was associated with microchannel fluid dynamics, we undertook computational fluid modeling. This indicated that fluid shear stress in the square cross-section channel varied with location along the channel wall, with shear stresses highest at the mid-wall and approaching zero toward the corners (Fig 1E).

These findings establish the generation of a small caliber micro-physiologic vessel system comprised of highly flow-responsive endothelial cells capable of physiologically remodeling the conduit.

### Intraluminal endothelial bridges can form and self-assemble in a flowing microchannel

We next sought evidence for intraluminal bridge formation. For this, confocal microscopy images of 24 perfusion-fixed, Alexa Fluor-phalloidin-stained microchannels were 3D- reconstructed, and the entire length of each channel was scrutinized. Remarkably, a total of 268 transluminal endothelial cell bridges were found (11.2±9.5 bridges per channel, range 0-43), with over 90% of channels containing at least one bridge.

All transluminal bridges emanated from and connected to a lining endothelial cell, and all contained F-actin. However, there was also considerable diversity in bridge morphology, with variations in bridge thickness, the presence and location of a cell nucleus, and whether the bridge was comprised of a single or multiple cells. Accordingly, we determined that bridges could be sub-typed. Type 1 bridges crossed the lumen as a thread-like strand, approximately 1 µm in diameter, with an F-actin signal that could be discontinuous (Type 1a, Fig. 2A). Somewhat thicker more cord-like connections (2-3 µm) were also seen (Type 1b, Fig. 2A, Movie S1). In either case, a type 1 bridge did not contain the cell nucleus. Type 2 bridges contained either part (type 2a) or all (type 2b) of the cell nucleus (Fig. 2A, Movie S2). Type 3 bridges were multicellular with multiple nuclei (Fig. 2A, Movie S3). Bridges could exist as a single transluminal structure or as part of an inter-connected network of bridges. A bridge network typically entailed a dominant, multicellular bridge with additional bridges emanating from it or the bridge-wall junction (Fig. 2D). Bridges were often tented at the wall-bridge junction, implicating contractile forces (Fig. 2A, e.g. Type 1b and 2a bridges).

**Figure 2.**
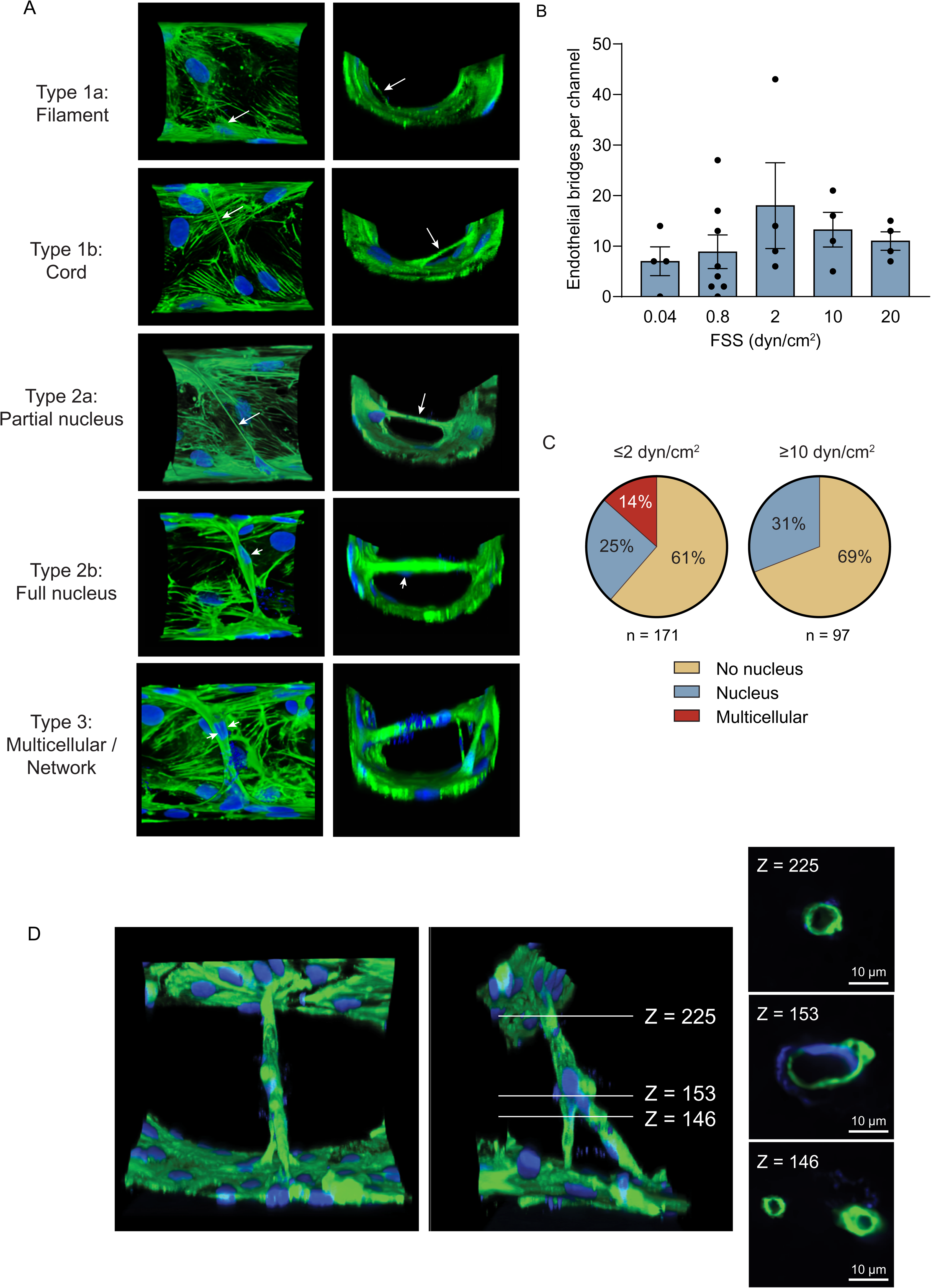
Endothelial cell bridges with diverse morphologies transect the lumen of micro-physiologic vessel conduit. **A**. Confocal volume projections of endothelialized microchannels constructed from z-slices (0.5 µm step size) showing transluminal endothelial bridges. Cells are stained for F-actin (Alexa Fluor-conjugated phalloidin, green) and nuclei (DAPI, blue). Each image pair depicts orthogonal views of different bridge morphologies, as labeled. Long arrows depict the bridging cell segment. Short arrows depict a cell nucleus either partly or entirely located within the bridge. In all bridges, cell tenting at the bridge-sidewall junction is evident, implicating contractile forces. Wall-to-wall maximum distance for all volume projections is 80 µm. See also Movie S1. **B.** Graph depicting the number of pillars identified per channel, for channels subjected to different flows. The corresponding fluid shear stresses are depicted. **C.** Chart showing the distribution of bridge morphologies observed under low and high shear stress conditions. **D.** Confocal volume projections of a complex, multicellular endothelial bridging structure spanning from top to bottom of the chamber. Two orthogonal views are shown as well as z-slice optical sections (0.5 µm step size) across the bridge. The structure can be seen to bifurcate, with one limb bifurcating again just above insertions into the channel wall. See also Movie S2. The F-actin and nuclear signals localize exclusively at the circumference of the structure with a signal void centrally, consistent with a self-organized structure that micro-divides the lumen up to 22 µm in the direction of flow. Z-slice numbers are shown.

Interestingly, bridges were identified in channels under all wall shear stress conditions tested, with no significant differences in the number of bridges (P=0.167, Fig. 2B). In contrast, bridge morphology was impacted by shear stress. Specifically, multicellular pillars were only observed under low shear stress conditions (<2 dyn/cm^2^). Type 1 bridges were the most prevalent in both low and high shear stress conditions (Fig. 2C, P=0.0001 and <0.0001, respectively).

Remarkably, reconstructed volume images revealed that some multicellular bridges had an identifiable core (Fig. 2E). Here, the F-actin signal and endothelial cell nuclei were localized exclusively at the bridge circumference, with a central signal void that could be up to 14 µm in the major axis (Fig. 2E). This advanced state of endothelial self-organization bore similarities to transluminal pillars and developing splits observed in histological sections (e.g. Fig. 7 in Patan et al. (35) and Figs. 1A and 2B in Dias-Flores et al (36)).

### Endothelial bridging within the microvessel-on-a-chip recapitulates *in vivo* bridging: Evidence from a systematic review of *in situ* intussusceptive angiogenesis

The diversity of bridge morphologies found in the microfluidic chamber raised the question as to whether this reflected transluminal bridges *in vivo*. However, many studies of IA have identified bridges and pillars through corrosion casting which does not inform on cellular structure. Even among histologic assessments, endothelial cell bridges are largely uncharacterized, relative to tissue pillars. Therefore, to understand the landscape of *in vivo* endothelial bridge morphologies, we undertook a systematic review of the literature to identify histological images of transluminal vascular bridges. PubMed structured searching for intussusceptive angiogenesis-based articles yielded 620 unique articles (*SI Appendix*, Fig. S1A). After review and application of exclusion criteria 148 articles were identified, all of which were scrutinized for images of transluminal bridges. This yielded 45 articles and 111 images of endothelial bridges (*SI Appendix*, Table S1). The bridges identified were from diverse contexts, including embryonic development, tumours, and inflamed, injured, or diseased adult tissues (*SI Appendix*, Fig. S1B). We ascertained that all bridges identified could be categorized as either: 1) a thin transluminal endothelial bridge without a nucleus, which corresponded to type 1 bridges observed in the microfluidic device; 2) a bridge containing part or all of a nucleus, corresponding to type 2 bridges, or 3) a multicellular bridge, which corresponded to type 3 bridges. The most common morphology was the thin bridge without a nucleus (*SI Appendix*, Fig. S1A), as also the case in the vessel-on-a-chip. Interestingly, five articles had images of vessels with a network of bridges, which could include bridges of any category, also a feature found in the microfluidic vessel model.

These findings reveal the current landscape of endothelial bridge morphologies and establish that the diversity identified was captured in the microfluidics-based micro-physiologic model vessel.

### Endothelial bridging within the microvessel-on-a-chip recapitulates *in vivo* bridging: Evidence from patients with limb-threatening ischemia

We previously found that IA was central to angiogenesis after ischemic hindlimb injury to the mouse (8). Therefore, we next prospectively examined the skin of lower limbs amputated from individuals with end-stage limb-threatening ischemia, an environment known to entail angiogenesis, although of uncertain mechanism (37, 38). Nine patients were studied, all of whom had ischemic skin wounds due to peripheral artery disease (PAD) (*SI Appendix*, Table S2). Specimens were immunostained for endoglin, an auxiliary receptor for TGF-ß that reliably identifies the microvasculature in ischemia-injured human skin, more so than CD31 (38).

In all patients, there was extensive skin vascularization, particularly in subdermal zones adjacent to the wound edge (Fig. 3A). Interestingly, double or twinned side-by-side capillaries were found, suggesting intussusceptive vessel duplication (Fig. 3B). Furthermore, microvessels with transluminal endothelial bridges were observed in all patients. The prevalence of bridges was greater at the wound edge than distal to the edge (n=7 tissues with paired comparisons, P=0.0302, Fig. 3C) (38). Moreover, a range of transluminal bridge morphologies was observed. This included thin strands (1 µm diameter), some of which were centrally located in the lumen (Fig. 3D) but others were eccentric and transected the lumen close to the vessel wall (Fig. 3E). There were also bridges in which an endothelial cell body and nucleus could be seen partially lining the wall and also extending into a lumen-transecting bridge (Fig. 3F). More mature bridges that fully contained one or more cell nuclei were also seen (Fig. 3G, H).

**Figure 3.**
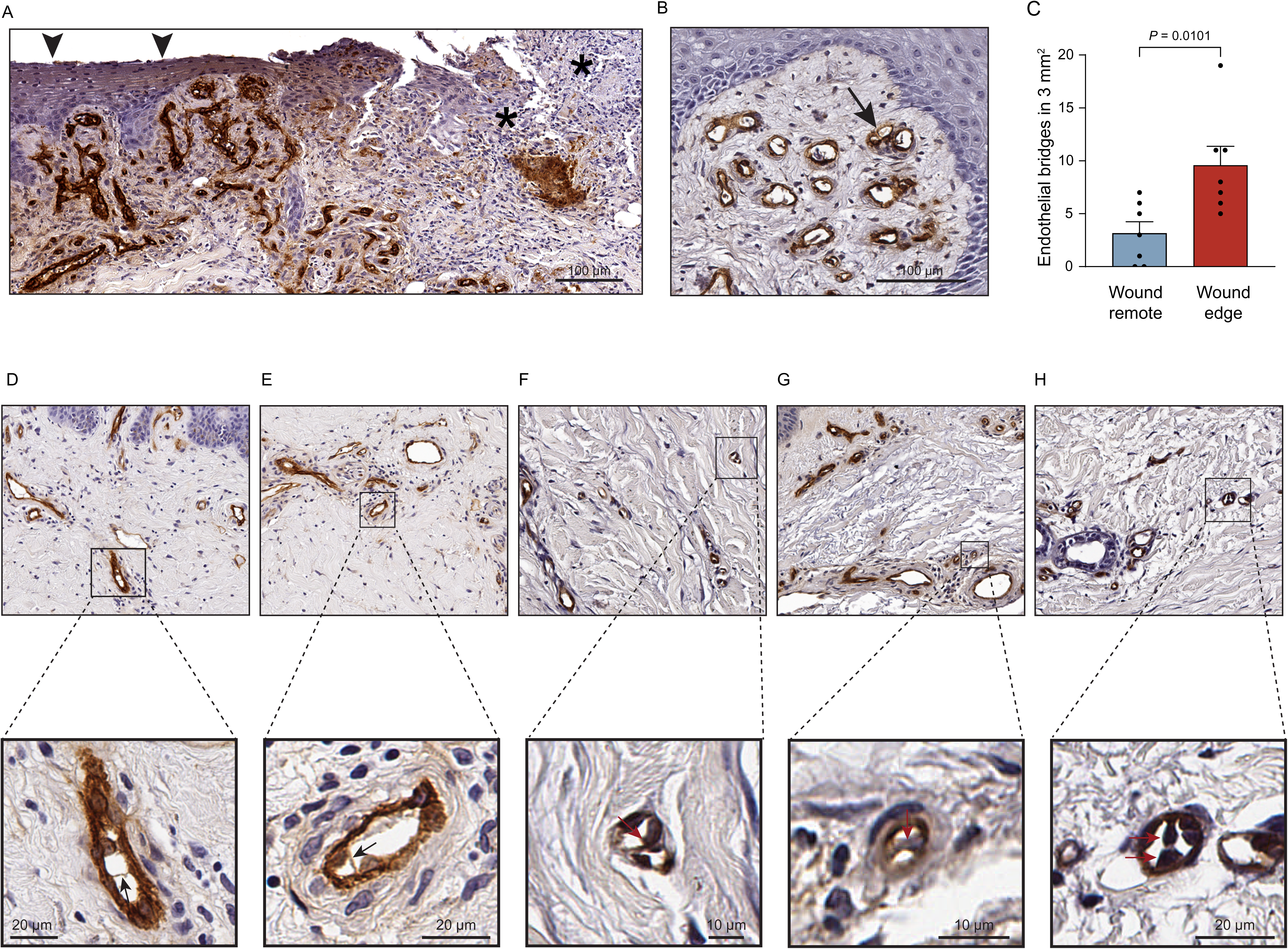
Transluminal endothelial bridging in microvessels in patients with limb-threatening ischemia. Sections of human skin from lower limbs amputated from of individuals with peripheral artery disease and non-healing skin wounds, immunostained for endoglin (brown). **A.** Image showing the chronic wound with avascular granulation tissue and absent epidermis (asterisk), and adjacent area with hyper-vascularized dermal zone (arrowhead). **B.** Section showing hyper-vascularized papillary dermal zone, including twinned microvessels (arrow). **C.** Graph depicting transluminal bridge content of microvessels adjacent to and remote (by >10 cm) from the wound. **D-H.** Low-magnification images, and zoomed images of microvessels within the corresponding boxes, illustrating: thin filament-like transluminal bridges (arrows, D, E); a bridge comprised of an endothelial cell and nucleus, part of which crosses the lumen and part of which lines the lumen (F); a thicker bridge of the same width as the microvessel wall and containing a cell nucleus (arrow, G); and a thicker bridge with two cell nuclei evident (arrows, H).

Thus, ischemic skin in patients with advanced PAD is an environment of transluminal bridging and IA. Diverse bridge morphologies existed in the microvessels of this tissue – morphologies that were recapitulated in the microfluidic model.

### Transluminal bridges form by partial delamination of circumferentially oriented endothelial cells

Having established close homology between *in vitro* and *in vivo* transluminal bridges, we next capitalized on the bridging phenomenon within the micro-physiologic conduit to investigate mechanistic processes. To understand how bridges formed, we undertook multi-plane 3D time-lapse confocal microscopy. Channels were lined with either GFP-expressing HUVECs or HUVECs incubated with LIVE 610-conjugated jasplakinolide and subjected to a shear stress of 1 dyn/cm^2^. Cells were imaged at sub-confluent cell densities to optimally delineate motility events, and images were acquired at 1 µm-steps through the channel height. This sequence was repeated every 10 minutes. Cells were color-mapped based on their vertical position.

Video analysis of height-mapped and volume-rendered images revealed surprisingly robust endothelial cell protrusive activity. From 283 dynamic cell protrusions, 42% protruded against the direction flow, 21% protruded in the direction of flow and, surprisingly, 37% protruded orthogonal to the direction of flow, i.e. circumferential protrusions. Notably, endothelial protrusion directly into the flowing lumen was not observed.

Remarkably however, we captured a process whereby circumferentially oriented endothelial cells partially lifted off the lumen wall. In such an occurrence, the atypically aligned cell would elongate, remain attached on either end to the wall or to an adjacent adherent endothelial cell and the intervening cell segment would delaminate from the wall to form a bridge. Coverage of the underlying wall was generally restored by adjacent cells. In some cells, the leading-edge protrusion and most of the cell body remained adherent, and the stretched trailing aspect of the cell delaminated to create the bridge (Fig. 4A, Movie S3). In other instances, delamination and bridge formation was biased toward the protruding cell front (Fig. 4B, Movie S4). We also observed bridges arising from two endothelial cells adherent to each other that circumferentially translocated in opposite directions (Fig. 4C, Movie S5). In another variation, a circumferentially oriented endothelial cell would release itself from an adjacent, circumferentially oriented cell and, with the ends still attached to the wall, snap into a bridging position (Fig. 4D, Movie S6).

**Figure 4.**
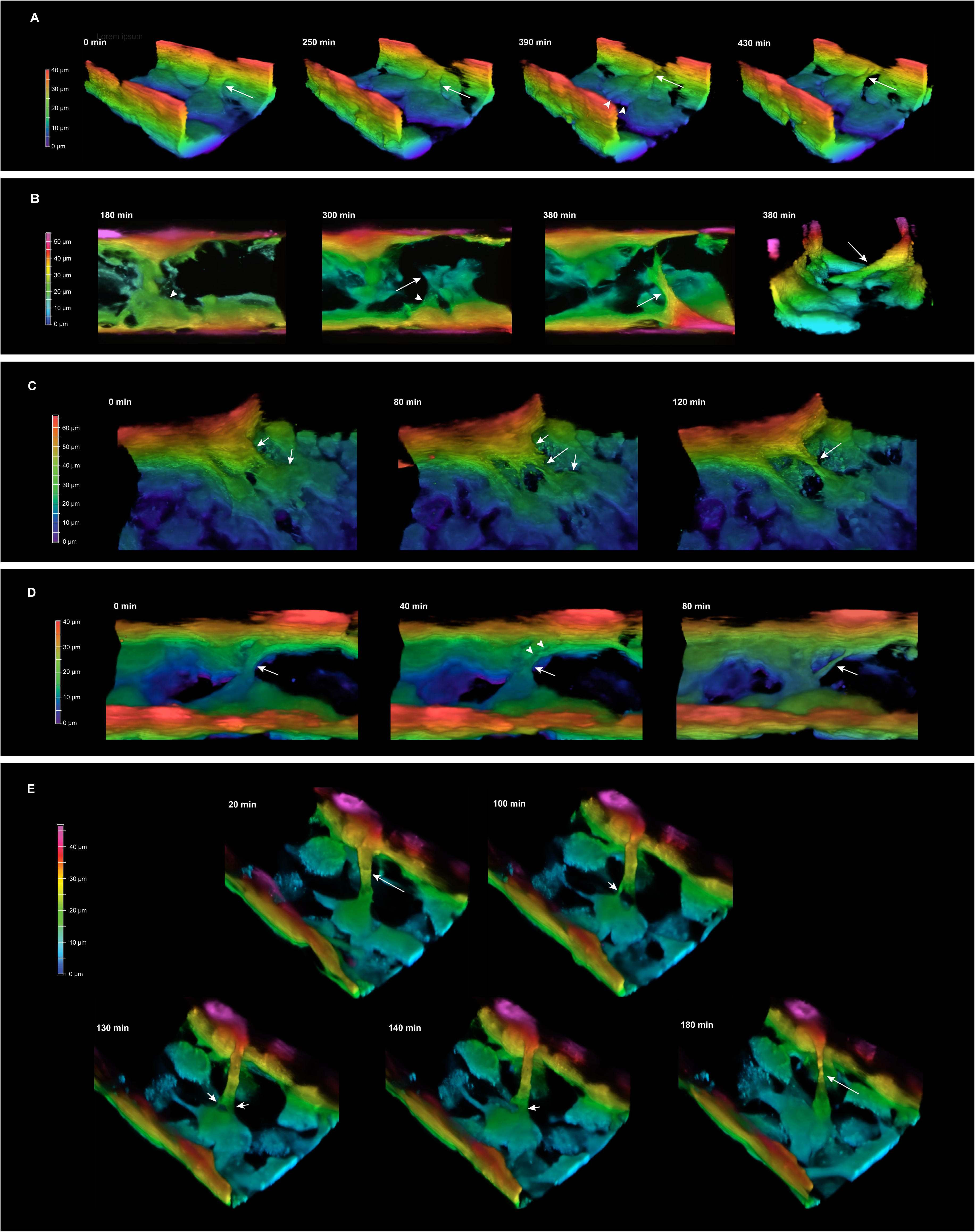
3D confocal time-lapse delineation of endothelial cell dynamics and transluminal bridging. **A.** Volume reconstructed 3D confocal time-lapse image frames of a GFP-expressing endothelial cell-lined microfluidic channel with cells subjected to 2 dyn/cm^2^ shear stress. Cells are color-mapped based on their vertical position within the channel. A circumferentially oriented endothelial cell is denoted by the arrows. A leading-edge protrusion (arrowheads, third panel) circumferentially crosses the channel bottom. This stretches the trailing aspect of the cell (third panel, arrow) which then partially delaminates with the distal tail remaining adherent, creating a cord-like bridge (fourth panel, arrow). See also Movie S3. **B**. 3D time-lapse image frames showing a GFP-expressing endothelial cell (arrowheads) crawling along the sidewall and then sending a long slender protrusion circumferentially along the channel floor (long arrow, panel 2) which partially delaminates to form a bridge (panel 3). The fourth panel shows an orthogonal reconstruction revealing the bridge architecture (arrow). See also Movie S4. **C.** 3D time-lapse image frames of endothelial cells labeled with the fluorescent actin probe, LIVE 610-conjugated jasplakinolide, showing two adjacent endothelial cells adhering to each other (short arrows). The cells circumferentially migrate apart from each other but remain connected (long arrow). The resulting stretched cell connection region releases from the wall to form a cord-like bridge (long arrow, panel 3). See also Movie S5. **D.** 3D time-lapse frames of GFP-expressing cells showing two cells extending circumferentially from bottom to lateral surfaces. In the first panel, the two cells are not discernible (arrow) but in the second panel a minor separation is evident (arrowheads). The upstream cell snaps away from the adjacent cell, retaining attachments on the bottom and lateral surfaces and the intervening segment forms a bridge (arrow, third panel). See also Movie S6**. E.** 3D time-lapse image frames of GFP-expressing endothelial cells showing a transluminal bridge (arrow) with one end of the bridging cell attached to the side wall and the other to an endothelial cell on the bottom surface. The bridge attenuates at its connection with the bottom cell (small arrow, panel 2) as the latter migrates. However, a second attachment with an adjacent bottom surface cell consolidates (panels 3 and 4, small arrow on the right) as the attachment to the index endothelial cell is released (panel 4), effectively passing the pillar from one cell to the adjacent once. The pillar itself (arrows, panels 1 and 5) lasted for 6 h. See also Movie S6. The distance from one wall of the channel to the opposite wall in the volume projections, evident in A, B, C, and E, is 80 µm.

Time-lapse 3D imaging also revealed that, once formed, pillars were dynamic. This could include rupturing of the bridge, but bridges could also reorganize their cell-cell connections. For example, a “bridge walking” phenomenon was observed wherein a cell process would attenuate and release from one site while forming a new adjacent attachment, effectively passing the bridge from cell to cell (Fig. 4E, Movie S7).

Collectively, these findings uncover previously unrecognized endothelial cell behaviors, the most striking being localized delamination from the ECM-coated conduit wall to form a dynamic bridge structure.

### Bridge entrance of endothelial cells into a flowing lumen depends on actomyosin tuning

We next investigated intracellular machinery that underlies bridge formation. To determine if bridging was actin-dependent, channels were infused with a low concentration of cytochalasin D. This disassembled the parallel microfilament bundles leaving only fragmented stress fibers and cortical actin (Fig. 5A). In association with these changes, there was an 85% decline in the number of transluminal bridges (P=0.0033, Fig. 5A), establishing the importance of the actin cytoskeleton in the bridging process.

**Figure 5.**
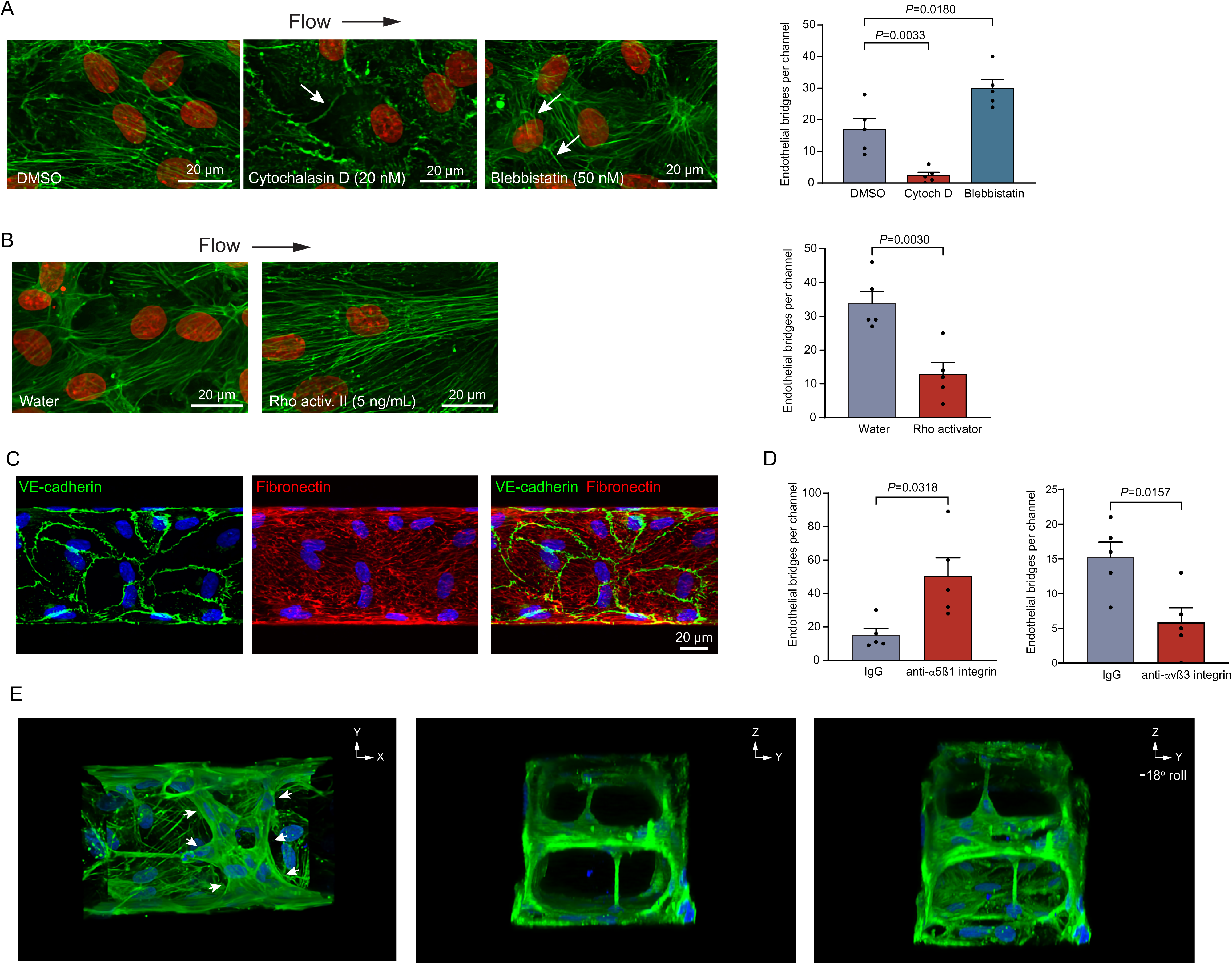
Role of the actin-myosin cytoskeleton and integrins in transluminal endothelial cell bridging. **A.** Maximum intensity projections of confocal fluorescent images of the bottom surface of an endothelial cell-lined microfluid chamber subjected to 2 dyn/cm^2^ shear stress. Cells are stained with Alexa Fluor-conjugated phalloidin (F-actin, green) and DAPI (nuclei, blue). Cytochalasin D resulted in loss and fragmentation of actin stress fibers and prominent and cortical actin (arrow). Blebbistatin resulted in shorter and less aligned actin stress fibers (arrows). The corresponding effects on transluminal bridge formation are shown on the right. **B.** Cells subjected to Rho kinase activation and imaged as in *A*, showing accentuated actin stress fibers and reduced transluminal bridge formation. **C.** Confocal fluorescent micrographs of the bottom surface of endothelial cell-lined microfluidic chamber subjected to 2 dyn**/**cm^2^ flow immunostained for VE cadherin (green) and nuclear-stained with DAPI (blue), with the underlying substrate immunostained for fibronectin fibrils (red) 72h after cell seeding. **D.** Graphs showing the effect of blocking α5ß1 integrin and αv3 integrin on transluminal bridge formation. **E.** Volume projections of 3D-reconstructed confocal images of an endothelialized microfluidic channel perfused with anti-α5ß1 integrin blocking antibody. The lumen is divided by a multicellular layer of bridging endothelial cells (arrows). The YZ and YZ rolled images further depict the neo-septum dividing the lumen into two parallel channels. Two additional bridges that connect the septum to opposing channel walls, transecting the two sub-lumens, are also present.

We also inhibited non-muscle myosin II (NMM-II) ATPase activity with blebbistatin, an intervention is known to relax actomyosin tension (39). This yielded shortened actin microfilament bundles (Fig. 5A) but not the overt stress fiber disassembly seen with cytochalasin D. Remarkably, this intervention led to a 76% *increase* in the number of transluminal bridges (P=0.0180, Fig. 5A).

We also infused channels with Rho activator II which induced assembly of thick, densely packed actin stress fibers (Fig. 5B). Interestingly, although a divergent actin response from cytochalasin D, this intervention also reduced the number of transluminal bridges, by 60% (P=0.0030, Fig. 5B).

Thus, transluminal bridging was inhibited by global actin disruption and by hyper-consolidating the microfilament bundles but facilitated when NMM-II-driven contractility was suppressed. Therefore, the shift from a lining to bridging position requires a functioning but relaxed state of the endothelial actomyosin machinery.

### Fibronectin-binding integrins have distinct roles in pillar formation

We next investigated the role of ECM receptors, recognizing their fundamental roles in adhesion, migration, and regulation of the cytoskeleton. Because the endothelial cells were seeded onto soluble fibronectin-coated PDMS, we first assessed if the endothelial cells assembled a fibronectin fibril network, a process that requires integrin engagement (40–42). Within 72 h of cell seeding, assembled fibronectin fibrils were immunodetectable on the basal endothelial surface (Fig. 5C). We thus next assessed the role of α5ß1 integrin in the bridging process by infusing α5ß1 integrin blocking antibody 24 hours after cell seeding. This led to a striking 230% increase in transluminal bridges (P=0.0318, Fig. 5D). Notably, this intervention also altered bridge morphology and promoted the formation of multicellular bridges. Such bridges not only transected the lumen but could be assembled as sheets running parallel to the direction of flow, effectively dividing the lumen (Fig. 5E).

We then tested the role of αvß3 integrin, another fibronectin-binding integrin on endothelial cells (43). Interestingly, infusion of the αvß3 integrin-blocking antibody, LM609, yielded a 62% *decrease* in the number of transluminal bridges (P=0.0157, Fig. 5D). LM609 is known to inhibit endothelial cell motility (44, 45) and this finding thus supports the importance of the observed migratory events in bridging.

Collectively, these findings establish multi-faceted roles for endothelial cell-fibronectin interactions, with differential participation of integrins in the transluminal bridging program.

## Discussion

Using a microfluidic vessel-on-a-chip strategy and 3D and 4D interrogation, we provide direct evidence that endothelial cells can autonomously leave their location lining the wall and transect a flowing lumen. They do so by a delamination process, lifting away from the wall while remaining viable and firmly attached at their two ends. The resulting bridges mimicked those found in several IA environments *in vivo*.

Generating a conduit with a small-caliber lumen circumferentially lined by healthy, flow-sensitive endothelial cells proved to be key to modeling early IA. Most cells in the conduit polarized against the direction of flow, a physiologic response that was evident even at the low shear stress of 0.8 dyn/cm^2^, albeit with more heterogeneity than at 10 dyn/cm^2^. Furthermore, the endothelialized conduit inwardly remodeled at zones of predicted ultra-low shear stress, yielding a more physiologic, rounded cross section. These vessel-like attributes are noteworthy because modeling fully-lined microvessels in a fluidic system is notoriously difficult (46, 47) due to the low cell seeding efficiency and the low media volume-surface area ratio, conditions that can compromise endothelial cell health. We overcame this using an interactive, visualization-based approach for cell seeding which greatly augmented seed coverage and thereby minimized the replicative aging stresses otherwise imposed as the cells divide to obtain confluence.

The ability to look inside the lumen at high resolution allowed us to discover a remarkable range of endothelial bridge morphologies. From 268 observations, we could classify bridges as: 1) strands or cords that did not contain the cell nucleus, 2) a single cell body including part or all of the nucleus, and 3) multicellular. Thus, diverse assemblies inside the lumen can emerge, all in an endothelial cell-autonomous manner. An especially remarkable level of endothelial cell self-organization was the finding of bridges comprised of multiple endothelial cells that surrounded a cell-free core. These structures resembled tissue pillars and early intraluminal divisions in vessels undergoing splitting *in vivo*. Pillars and early splits have a core that can include myofibroblasts and collagen (5, 48), and ascertaining if such a program can be modeled *in vitro* will require multi-component micro-physiologic system innovations. Nonetheless, the current findings raise the possibility that lumen-transecting endothelial cells can autonomously reorganize within a flowing lumen to facilitate the influx of mesenchymal cells as part of the vessel duplication process.

The finding of diverse endothelial assemblies within the vessel-on-a-chip stimulated us to better understand the structural features of endothelial bridges *in vivo*, which has not been well characterized. The systematic review of published transluminal bridge images – covering a range of species, tissue beds, developmental stages, and normal and diseased contexts – provided a synthesis of cell details. Interestingly, the correspondence between the self-organized structures in the vessel-on-a-chip and those found *in vivo* applied not only to their morphologies but also their relative abundance.

We also found transluminal endothelial bridges in dermal zones adjacent to ischemic wounds in patients with limb-threatening ischemia. This is important for several reasons. First, the close homology to bridge morphologies *in vivo* was again demonstrated. Second, there is a pressing need to better understand microvascular structure and function in PAD. Advanced PAD with non-healing ulcers, including in diabetics, is a debilitating ischemic disease that carries a high risk for limb loss or death (49). Therapeutic strategies to stimulate sprouting angiogenesis in this condition have not been successful (50, 51). We speculate that most of the bridge morphologies that we identified in this diseased tissue constituted an early phase of IA, supported by the presence of double or twinned capillary profiles which also implicate IA (52). A predominance of IA in advanced PAD could in part explain the failure of strategies that, instead, were designed to promote sprouting angiogenesis. It is also possible that some of the transluminal bridges in the dermal microvessels are pathologic structures that might prevent optimal transit of red cells, a phenomenon that has been identified in vascular malformations (15). Also noteworthy is that some of the transluminal bridges in the PAD microvessels were asymmetrically positioned very close to the vessel side-wall, a localization also identified in ischemic mouse skeletal muscle (8). Although inferential, this *in situ* observation is consistent with a delamination event.

The phenomenon of transluminal endothelial bridges arising from controlled delamination from the wall was directly supported by 3D confocal time-lapse imaging. The detachment process was partial, with the cell ends remaining adhered to the substrate and adjacent endothelial cells. This process within the 3D and fluidic environment is strikingly different from the tail deadhesion and retraction phenomenon of cells migrating on planar surfaces (53). We also found no instances of cells rounding and lifting off entirely, or other evidence for anoikis. Importantly, although most endothelial cells in the channel were oriented against or with the direction of flow, bridging was identified exclusively in circumferentially oriented cells. Elongation of such a cell meant acquiring a uniquely bent conformation, corresponding to the lumen curvature. We propose that a circumferentially oriented endothelial cell and the resulting bent morphology is a vital set-up for converting an otherwise adherent endothelial cell to a bridge.

Several other mechanistic determinants of transluminal bridging were established. Bridging was actin-dependent, consistent with the need for dynamic positioning of the bridge precursor cell. The requirement for cell repositioning was supported by the observed suppression of bridging upon over-activation of Rho. The resulting fortified stress fibers and associated augmented adhesion to the substrate would be expected to limit both dynamic cell repositioning and partial release from the substrate. On the other hand, inhibiting NMM2 without obliterating stress fibers is an intervention that could facilitate motility and partial de-adhesion, consistent with the enhanced bridging observed with this intervention. Hence an intact but tunable and relatively relaxed state of actomyosin is required for the transition of an endothelial cell from wall to lumen. The actin dynamics required for lining endothelial cells to become bridges are undoubtedly complex and likely differ from the dynamics in cells already bridging the lumen. Within more well-developed bridges, there was evidence for robust contractile forces, with thick actin stress fiber bundles along the length of the bridge and tenting of cells at the bridge-sidewall interface. These forces might draw additional endothelial cells into the bridge, or as additional bridges.

The paradigm of bridging by delamination was further supported by the enhanced bridge formation upon blocking α5ß1 integrin. Not only did more bridges form with this intervention but bridges were more complex, including thick, multicellular transecting structures and endothelial sheets that effectively split the lumen into two parallel lumens. Interestingly, blocking αvß3 integrin, another fibronectin-binding integrin, suppressed bridging. LM609 is known to preferentially inhibit endothelial cell motility (44, 45). This finding is thus consistent with the need for endothelial cells to actively position themselves, including elongating along the curvature, as a prerequisite for bridging.

We have previously found that low shear stress and reduced shear sensing competence are enabling conditions for bridge or pillar formation in skeletal muscle neovessels (8). The current findings indicate that the magnitude of bulk fluid shear stress is not, by itself, a dominant driver of the decision to form a bridge. However regional heterogeneity in shear stress, which has been proposed to underlie IA, still could be (54). Shear stress modeling of the microfluidic chamber established that there can be profound local differences in shear stress depending on wall geometry. As well, the finding that ultra-low shear stress favoured complex bridge formation also supports a functional interplay between shear stress conditions and endothelial bridge assembly.

At a conceptual level, the phenomenon of an endothelial cell entering and crossing a flowing lumen seemingly defies the principles of substrate-dependent cell movement. Our discovery that a bridge can arise through delamination offers a resolution to this paradox. We propose a model of transluminal endothelial cell bridging (Fig. 6) that begins with elongation of one or more endothelial cells that are polarized orthogonal to the direction of flow. This is followed by local de-adhesion and lifting of the bent cell segment away from the vessel wall. The integrity of the underlying wall could be preserved through spreading of adjacent endothelial cells or if the presumptive lifting cell initially overlapped with an adjacent endothelial cell (7). The newly formed bridge can rupture or can consolidate and develop by recruiting additional endothelial cells from the wall and by repositioning bridge endothelial cells including to surround an endothelial cell-free space.

**Figure 6.**
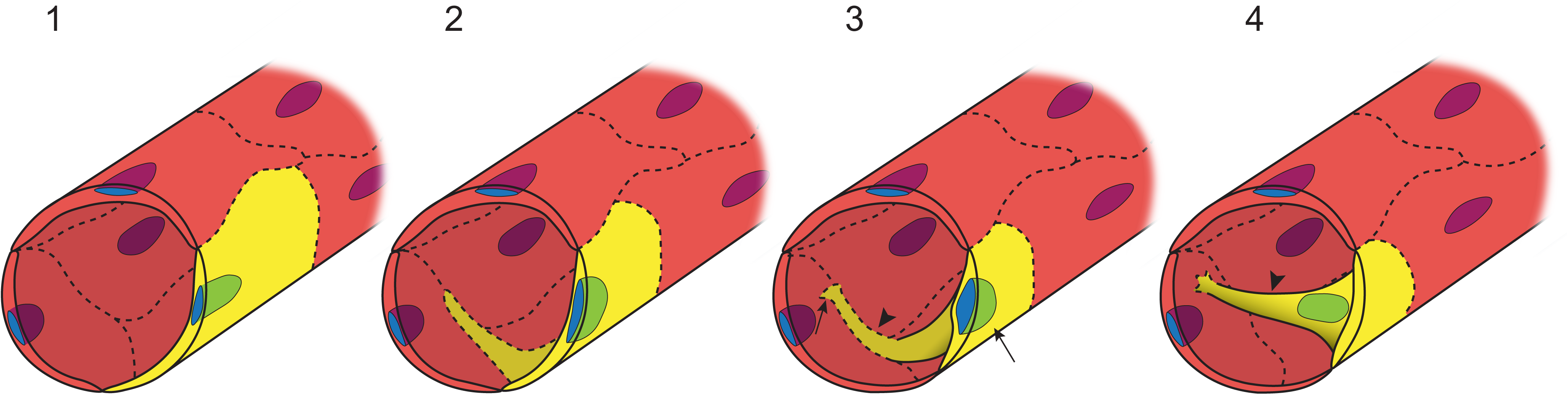
Schematic depiction of transluminal endothelial bridge formation in microvessels. Endothelial cells co-align and axially orient by mechanosensing flow (*1*). A presumptive bridging cell is shown in yellow and does not sustain this orientation but repositions and circumferentially extends a cytoplasmic process (*2*). The leading edge of the protrusion and the trailing cell body (arrows) remain adherent and within the monolayer, but the intervening component delaminates from the substrate (arrowhead) (*3*). This liberates the circumferentially oriented process to form a transluminal bridge (*4*). The repositioning and selective delamination processes may arise from an interplay between maintained αvß3 integrin activity and reduced α5ß1 integrin activity. Cytoskeletal control is dynamic and may draw the nucleus into the bridge.

## Limitations

Because the microfluidic vessel was not of capillary dimensions we could not investigate the previously proposed paradigm of opposing endothelial cell surfaces contacting and reorganizing into a bridge (48). Capillaries have yet to be modeled with endothelialized microfluidic channels. Nevertheless, the delamination mechanism we identified could conceivably proceed in capillaries, in addition to larger microvessels where opposing cell surfaces cannot feasibly contact each other. As well, although we did not identify evidence for endothelial protrusions entering and crossing the flowing lumen, we cannot exclude the possibility. Higher temporal resolution might capture such activity. The core components of the pillar-like structures that formed in the micro-physiologic conduit system are currently unknown. Histological sectioning of PDMS is not productive but advances in multiplex *in situ* labelling and multi-component micro-physiologic models could yield exciting data in this regard (55). Finally, a key benefit of our model is that it enables imaging of rapid and transient single-cell events inside a vessel lumen, but similar advances will be required to investigate endothelial delamination *in vivo*.

## Summary

We report the first live-cell model of transluminal endothelial cell bridging. The interrogation enabled by this model revealed that select endothelial cells can leave their monolayer to enter and transect the flowing lumen through controlled delamination. The resulting dynamic bridges had morphologies homologous to bridges and pillars *in vivo*, including in patients with limb-threatening ischemia. Thus, in addition to serving as a model system for understanding the early events of IA, the micro-physiologic system herein could serve as a platform for seeking interventions that promote or inhibit non-sprouting angiogenesis.

## Materials and Methods

### Microfluidic device fabrication

A custom designed microfluidic device was assembled using a previously reported strategy (56). Briefly, a replica mold, 80 µm x 80 µm x 1 cm, was machined on the end surface of an acrylic cylinder using a computer numerical control milling machine. Acrylic collars, 2.65 mm in diameter and 1.56 mm in height, were pinned and glued to the ends of the channel mold and served as molds for inlet and outlet reservoirs. The complete mold was mounted in a custom-milled acrylic sleeve and covered with 1.5 mL of vacuum-degassed polydimethylsiloxane (PDMS, Sylgard 184; Dow Corning). The PDMS was cured at 60°C for 1 hour after which the cast imprinted with the channel and inlet/outlet reservoirs was removed. The channel was closed by generating the bottom surface. PDMS (200 µL) was applied to the center of a glass-bottomed cell culture dish (MatTek 50 mm, 30-mm glass bottom) and spun for 60s at 3000 RPM to produce a thin coating. The coating was partially cured at 60°C and the coated dish irreversibly attached to the PDMS cast by gentle pressure and curing for 20 minutes at 60°C. PE50 tubing (Becton-Dickinson) was inserted into the pin-derived holes within the inlet and outlet and secured with epoxy.

### Culture of endothelial cells in microchannels

Prior to cell seeding, microchannels were infused with fibronectin (250 µg/ml; F1141, Sigma-Aldrich) at 1 µL/min for 1 hour using a programmable syringe pump (NE-300 or NE- 1002X, New Era Pumps). Channels were then infused with endothelial growth media 2 (EGM-2, CC-3162, Lonza) supplemented with 0.1 µg/mL L-ascorbic acid (A4034, Sigma-Aldrich) at 1 µL/min for 30 minutes. Human umbilical vein endothelial cells (HUVECs, CC-2519, Lonza, passage 4-7) cultured in EGM-2 were then trypsinized, resuspended at 1 x 10^6^ cell/mL in ascorbic acid-supplemented EGM-2 containing 8% dextran, and perfused into the microchannels under direct visualization using Hoffman-modulated contrast microscopy (Zeiss, Axiovert S100). During this time, cell delivery into the channel was tightly regulated by adjusting the heights of the delivery and exit tubing. With this, flow could be stopped and slowly restarted as appropriate while visualizing cell attachment to the channel walls, which optimized the cell attachment process. Once maximum coverage was achieved, the cell-lined channels were incubated for 15-18 hours without flow to avoid shear stress-induced suppression of proliferation. Endothelial cells were then acclimatized to shear stress by perfusing channels at 1 µL/min for 24 hours. Channels were then perfused at 0.02, 0.4, 1, 5 or 10 µL/min for 48 hours, corresponding to applied shear stresses of 0.04, 0.8, 2, 10 and 20 dyn/cm^2^, respectively. Shear stress was estimated using the Navier-Stokes and continuity equation-derived formula, τ = 6Qµ/(bh^2^), where τ is shear stress, Q is flow rate, µ is viscosity, and b and h are channel width and height, respectively. The viscosity of culture media was assumed to be that of water (1 cP).

### Immunofluorescent staining of endothelial cell-lined microchannels

Cells were perfusion-fixed *in situ* with 4% paraformaldehyde (1 µL/min x 10 min), permeabilized with 0.1% Triton-X in phosphate buffer solution (PBS, 1 µL/min x 10 min) and blocked with 5% BSA + 0.1% Tween 20 in PBS (1 µL/min x 1 h). Antibodies were diluted in blocking solution and infused at 1 µL/min. Goat polyclonal anti-VE-cadherin antibody (1:100; R&D Systems, AF1002, 1 µL/min x 1h) was visualized with Alexa Fluor 488–conjugated donkey anti-goat IgG (1:1000; Thermo Fisher Scientific, 1 µL/min x 1h). Mouse monoclonal antibody anti-Golgi matrix protein 130 (GM130, 1:200; BD Biosciences, 1 µL/min x 1h) was visualized with Alexa Fluor 546-conjugated donkey anti-mouse IgG (1:500; Invitrogen, 1 µL/min x 1h). The F-actin cytoskeleton was visualized with Alexa Fluor 549 or Alexa Fluor 647 conjugated phalloidin (1 µL/min x 30 min). Nuclei were visualized with either DR (1 µL/min x 15 min) (1:200; Thermo Fisher Scientific) or 4′,6-diamidino-2- phenylindole (DAPI) in Fluoromount-G mounting medium (1 µL/min x 30 min, SouthernBiotech, 0100-01).

### Cytoskeletal interventions

Cytoskeletal modifying reagents employed were cytochalasin D (C8273, Sigma-Aldrich): 20 nM, Rho activator II (CN03, Cytoskeleton): 5 ng/mL, blebbistatin (B0560, Sigma-Aldrich): 50 nM, mouse monoclonal anti-α5β1 (JBS5 clone, MAB1969-I, Sigma-Aldrich), anti-αvβ3 (LM609 clone, ab190147, abcam) and IgG1, kappa control (ab170190, abcam). Reagents were added to culture medium at the onset of flow, 24 hours after cell seeding. All antibodies were diluted to 0.5 µg/mL.

### Confocal microscopy imaging of endothelial cell-lined microfluidic channels

Endothelial cell-lined channels were imaged using a Nikon Ti2-E Inverted Microscope and A1R HD laser-scanning confocal system with 405, 488, 561 and 640 nm lasers. A 20x water-immersion objective with numerical aperture of 0.95 and a 2x digital zoom was used to acquire z-stacks (80-220 slices) with a 0.5 µm z-step size. Confocal images were 12-bit (1024×512 pixels), two scan-averaged with an X-Y resolution of 620 nm. 3D volume projections were generated from either lower half or full-height acquisitions using the alpha blending mode in NIS Elements software (Nikon).

### Time lapse imaging of endothelial cell translocation activity in microfluidic channels

To visualize endothelial cell locomotion within the device, microchannels were seeded with GFP-expressing HUVECs (Angio-Proteomie, cAP-0001GFP) or, alternatively, with HUVECs exposed for 18 h to LIVE 610-conjugated jasplakinolide (0.01 µM, Abberior, LV610). For either method, 16 hours prior to imaging, cell culture media was supplemented with 1 µg/mL L-ascorbic acid to reduce phototoxic effects. HUVECs were subjected to flow corresponding to 2 dyn/cm^2^ for 40 hours then transferred to a humidified live-cell imaging chamber at 37℃ with 5% CO_2_, 21% O_2_ and 74% N_2_. Wall shear stress was maintained at 2 dyn/cm^2^ for the duration of imaging. Live cells were imaged with a Nikon Ti2-E Inverted Microscope and A1R HD laser-scanning confocal system using a 20x objective with 2x digital zoom and 488 or 640 nm lasers. Live imaging was undertaken in extended regions of interest at each end of the channel and 3D volumes acquired using a 1 µm step size. Each acquisition entailed four adjacent images of the same plane stitched together with 15% overlap (total area, 1137 µm x 160 µm), using ND acquisition software (Nikon). Images were captured every 10 minutes for 8 hours.

### Endothelial cell polarity analysis

Endothelial cell polarization was assessed based on positioning of the Golgi apparatus, ascertained in maximum intensity projections using ImageJ. A polarization angle was determined based on a line connecting the center of the Golgi apparatus with the center of the cell nucleus and a line parallel to the direction of flow. Cells with polarization angles ≤ 60° were classified as oriented with flow, with angles ≥ 120° as oriented against flow, and with angles between 60° and 120° as orthogonal to flow.

### Systematic review of transluminal endothelial bridge images

A search for all publications related to intussusceptive angiogenesis was performed using Medline database from its inception in 1946 to July 4th, 2024. Databases with earlier inceptions were considered unnecessary as the earliest published images depicting intussusceptive angiogenesis were in 1986. The search terms were: {intussusceptive angiogenesis}, {intussusceptive capillary growth}, {intussusceptive microvascular growth}, {splitting angiogenesis}, {pillar angiogenesis} and {“luminal division”}. The resulting records, including titles and abstracts were imported into the online systematic review management software Covidence, and duplicated records removed. All records were screened to verify being a study of intussusceptive angiogenesis based on title and abstract. Review articles, editorials, non-English language publications, and studies of intussusceptive lymphangiogenesis were excluded. After exclusions, the full texts of 148 articles were examined for the presence of histological images of nucleus-stained vessel cross-sections reported to depict IA. Studies that identified pillars based only by visualizing the intravascular space without cell staining or labeling, as in corrosion casting, were not included, nor were studies with only indirect evidence of IA (e.g. differentially expressed genes associated with IA).

The final set of studies included 45 English language, peer-reviewed manuscripts of original research containing cross-sectional histological images of transluminal endothelial bridges. For each image, the organism, physiological context, number of bridges, and morphology of bridges were ascertained (*SI Appendix*, Table S1).

### Dermal microvessel analysis in limb-threatening ischemia

Human skin samples from nine individuals (eight males, one female) who underwent lower limb amputation because of end-stage limb ischemia and chronic non-healing skin wounds were studied. Seven of the subjects had type 2 diabetes. Skin samples, described in a prior report (38), were fixed with 4% paraformaldehyde, embedded in paraffin, and 5-µm-thick sections were immunostained for endoglin (anti-endoglin/CD105, sc-376381, Santa Cruz Biotechnology) with DAB peroxidase colorimetric development. For each patient, a section at the wound edge, ie adjacent to the zone with absent epidermis, and a section at least 10 cm away from the wound were studied. Sections were scanned using a Leica Aperio AT2 bright field digital slide scanner with 40× objective, and microvessels of the superficial dermis within a 1 × 3 mm region of interest were quantified and scrutinized for transluminal bridges. Bridges in open-lumen capillaries and arterioles were counted and expressed per unit area, recognizing that quantifying microvessel density was confounded by the irregular angiogenesis network with many collapsed lumens.

## Statistics

Data from distributions passing Shapiro-Wilks normality test are presented as mean ± standard deviation (SD) for descriptive data and mean ± standard error of the mean (SEM) for compared data. For normally distributed data, comparisons were made by Student’s t-test or one-way analysis of variance (ANOVA) with Tukey’s post-hoc test. Differences in the distributions of endothelial cell polarization under different shear stress conditions were assessed using a linear mixed model with post-hoc pairwise comparison of marginal means. Differences between angiogenesis adjacent to and remote from the skin wound were undertaken by paired t-test. All statistical tests were two sided, and significance was set at P<0.05.

## Acknowledgements and Funding

This work was supported by the Canadian Institutes of Health Research (FDN-143326 and PJT-180604, J.G.P.), the Natural Sciences and Engineering Research Council of Canada (RGPIN-2020-05646, J.G.P.), and the Academic Medical Organization of Southwestern Ontario (INN22- 009, J.G.P.). J.G.P. holds the Neil McKenzie Chair in Cardiac Care at the Schulich School of Medicine and Dentistry, Western University.

## Data availability

The data underlying the findings are available in the article and supporting information.

## References

1. P. Carmeliet, R. K. Jain, Molecular mechanisms and clinical applications of angiogenesis. Nature 473, 298–307 (2011).

2. H. Yin et al., Fibroblast growth factor 9 imparts hierarchy and vasoreactivity to the microcirculation of renal tumors and suppresses metastases. J. Biol. Chem. 290, 22127–22142 (2015).

3. G. Eelen, L. Treps, X. Li, P. Carmeliet, Basic and Therapeutic Aspects of Angiogenesis Updated. Circ. Res. 127, 310–329 (2020).

4. J. H. Caduff, L. C. Fischer, P. H. Burri, Scanning electron microscope study of the developing microvasculature in the postnatal rat lung. Anat. Rec. 216, 154–164 (1986).

5. P. H. Burri, M. R. Tarek, A novel mechanism of capillary growth in the rat pulmonary microcirculation. Anat. Rec. 228, 35–45 (1990).

6. S. Patan, M. J. Alvarez, J. C. Schittny, P. H. Burri, Intussusceptive microvascular growth: a common alternative to capillary sprouting. Arch. Histol. Cytol. 55 Suppl, 65-75 (1992).

7. S. Egginton, A. L. Zhou, M. D. Brown, O. Hudlicka, Unorthodox angiogenesis in skeletal muscle. Cardiovasc. Res. 49, 634–646 (2001).

8. J. M. Arpino et al., Low-flow intussusception and metastable VEGFR2 signaling launch angiogenesis in ischemic muscle. Sci Adv 7, eabg9509 (2021).

9. D. Ribatti et al., Microvascular density, vascular endothelial growth factor immunoreactivity in tumor cells, vessel diameter and intussusceptive microvascular growth in primary melanoma. Oncol. Rep. 14, 81–84 (2005).

10. R. Hlushchuk et al., Tumor recovery by angiogenic switch from sprouting to intussusceptive angiogenesis after treatment with PTK787/ZK222584 or ionizing radiation. Am. J. Pathol. 173, 1173–1185 (2008).

11. B. Nico et al., Intussusceptive microvascular growth in human glioma. Clin Exp Med 10, 93–98 (2010).

12. D. Ribatti, V. Djonov, Intussusceptive microvascular growth in tumors. Cancer Lett. 316, 126–131 (2012).

13. M. A. Konerding et al., Inflammation-induced intussusceptive angiogenesis in murine colitis. Anat Rec (Hoboken) 293, 849–857 (2010).

14. M. Ackermann et al., Pulmonary Vascular Endothelialitis, Thrombosis, and Angiogenesis in Covid-19. N. Engl. J. Med. 383, 120–128 (2020).

15. W. Li et al., Abortive intussusceptive angiogenesis causes multi-cavernous vascular malformations. eLife 10 (2021).

16. M. Reichardt et al., 3D virtual histopathology of cardiac tissue from Covid-19 patients based on phase-contrast X-ray tomography. eLife 10 (2021).

17. C. Werlein et al., Inflammation and vascular remodeling in COVID-19 hearts. Angiogenesis 26, 233–248 (2023).

18. V. Djonov, M. Schmid, S. A. Tschanz, P. H. Burri, Intussusceptive angiogenesis: its role in embryonic vascular network formation. Circ. Res. 86, 286–292 (2000).

19. S. Patan, B. Haenni, P. H. Burri, Implementation of intussusceptive microvascular growth in the chicken chorioallantoic membrane (CAM): 1. pillar formation by folding of the capillary wall. Microvasc. Res. 51, 80–98 (1996).

20. H. Ross, D. Aaldijk, M. Vladymyrov, A. Odriozola, V. Djonov, Transluminal Pillars-Their Origin and Role in the Remodelling of the Zebrafish Caudal Vein Plexus. International journal of molecular sciences 24 (2023).

21. R. Gianni-Barrera et al., VEGF over-expression in skeletal muscle induces angiogenesis by intussusception rather than sprouting. Angiogenesis 16, 123–136 (2013).

22. R. Gianni-Barrera et al., PDGF-BB regulates splitting angiogenesis in skeletal muscle by limiting VEGF-induced endothelial proliferation. Angiogenesis 21, 883–900 (2018).

23. S. Paku et al., A new mechanism for pillar formation during tumor-induced intussusceptive angiogenesis: inverse sprouting. Am. J. Pathol. 179, 1573–1585 (2011).

24. S. J. Mentzer, M. A. Konerding, Intussusceptive angiogenesis: expansion and remodeling of microvascular networks. Angiogenesis 17, 499–509 (2014).

25. C. Du Cheyne, M. Smeets, W. De Spiegelaere, Techniques used to assess intussusceptive angiogenesis: A systematic review. Dev. Dyn. 250, 1704–1716 (2021).

26. L. Jakobsson et al., Endothelial cells dynamically compete for the tip cell position during angiogenic sprouting. Nat Cell Biol 12, 943–953 (2010).

27. D. H. Nguyen et al., Biomimetic model to reconstitute angiogenic sprouting morphogenesis in vitro. Proc. Natl. Acad. Sci. U. S. A. 110, 6712–6717 (2013).

28. S. Lee et al., Angiogenesis-on-a-chip coupled with single-cell RNA sequencing reveals spatially differential activations of autophagy along angiogenic sprouts. Nature communications 15, 230 (2024).

29. M. S. Weiss, G. Trapani, H. Long, B. Trappmann, Matrix Resistance Toward Proteolytic Cleavage Controls Contractility-Dependent Migration Modes During Angiogenic Sprouting. Adv Sci (Weinh) 11, e2305947 (2024).

30. M. W. Laschke, Y. Gu, M. D. Menger, Replacement in angiogenesis research: Studying mechanisms of blood vessel development by animal-free in vitro, in vivo and in silico approaches. Frontiers in physiology 13, 981161 (2022).

31. R. M. Hirschberg, M. Sachtleben, J. Plendl, Electron microscopy of cultured angiogenic endothelial cells. Microsc. Res. Tech. 67, 248–259 (2005).

32. A. M. Malek, S. L. Alper, S. Izumo, Hemodynamic shear stress and its role in atherosclerosis. JAMA 282, 2035–2042 (1999).

33. C. A. Franco et al., Dynamic endothelial cell rearrangements drive developmental vessel regression. PLoS biology 13, e1002125 (2015).

34. A. C. Vion et al., Endothelial Cell Orientation and Polarity Are Controlled by Shear Stress and VEGF Through Distinct Signaling Pathways. Frontiers in physiology 11, 623769 (2020).

35. S. Patan, B. Haenni, P. H. Burri, Evidence for intussusceptive capillary growth in the chicken chorio-allantoic membrane (CAM). Anat. Embryol. (Berl). 187, 121–130 (1993).

36. L. Diaz-Flores et al., Intussusceptive angiogenesis and its counterpart intussusceptive lymphangiogenesis. Histol. Histopathol. 35, 1083–1103 (2020).

37. J. Chevalier et al., Obstruction of small arterioles in patients with critical limb ischemia due to partial endothelial-to-mesenchymal transition. iScience 23, 101251 (2020).

38. J. Wang et al., Microcirculation surrounding end-stage human chronic skin wounds is associated with endoglin/CD146/ALK-1 expression, endothelial cell proliferation and an absence of p16(Ink4a). Wound Repair Regen. 31, 321–337 (2023).

39. C. Wilson, N. Naber, E. Pate, R. Cooke, The myosin inhibitor blebbistatin stabilizes the super-relaxed state in skeletal muscle. Biophys. J. 107, 1637–1646 (2014).

40. C. Wu, J. S. Bauer, R. L. Juliano, J. A. McDonald, The alpha 5 beta 1 integrin fibronectin receptor, but not the alpha 5 cytoplasmic domain, functions in an early and essential step in fibronectin matrix assembly. J. Biol. Chem. 268, 21883–21888 (1993).

41. S. Li, C. Van Den Diepstraten, S. J. D’Souza, B. M. Chan, J. G. Pickering, Vascular smooth muscle cells orchestrate the assembly of type I collagen via alpha2beta1 integrin, RhoA, and fibronectin polymerization. Am. J. Pathol. 163, 1045–1056 (2003).

42. J. G. Pickering et al., alpha5beta1 integrin expression and luminal edge fibronectin matrix assembly by smooth muscle cells after arterial injury. Am. J. Pathol. 156, 453–465 (2000).

43. D. A. Cheresh, Human endothelial cells synthesize and express an Arg-Gly-Asp-directed adhesion receptor involved in attachment to fibrinogen and von Willebrand factor. Proc. Natl. Acad. Sci. U. S. A. 84, 6471–6475 (1987).

44. D. I. Leavesley, M. A. Schwartz, M. Rosenfeld, D. A. Cheresh, Integrin beta 1- and beta 3-mediated endothelial cell migration is triggered through distinct signaling mechanisms. J. Cell Biol. 121, 163–170 (1993).

45. C. J. Drake, D. A. Cheresh, C. D. Little, An antagonist of integrin alpha v beta 3 prevents maturation of blood vessels during embryonic neovascularization. J. Cell Sci. 108 (Pt 7), 2655–2661 (1995).

46. M. B. Esch, D. J. Post, M. L. Shuler, T. Stokol, Characterization of in vitro endothelial linings grown within microfluidic channels. Tissue Eng Part A 17, 2965–2971 (2011).

47. Y. Zheng et al., In vitro microvessels for the study of angiogenesis and thrombosis. Proc. Natl. Acad. Sci. U. S. A. 109, 9342–9347 (2012).

48. P. H. Burri, R. Hlushchuk, V. Djonov, Intussusceptive angiogenesis: its emergence, its characteristics, and its significance. Dev. Dyn. 231, 474–488 (2004).

49. D. G. Armstrong, A. J. M. Boulton, S. A. Bus, Diabetic Foot Ulcers and Their Recurrence. N. Engl. J. Med. 376, 2367–2375 (2017).

50. B. H. Annex, Therapeutic angiogenesis for critical limb ischaemia. Nature reviews. Cardiology 10, 387–396 (2013).

51. S. R. Iyer, B. H. Annex, Therapeutic Angiogenesis for Peripheral Artery Disease: Lessons Learned in Translational Science. JACC. Basic to translational science 2, 503–512 (2017).

52. D. M. Dane et al., Inhalational delivery of induced pluripotent stem cell secretome improves postpneumonectomy lung structure and function. J Appl Physiol (1985) 129, 1051–1061 (2020).

53. S. Li, L. H. Chow, J. G. Pickering, Cell surface-bound collagenase-1 and focal substrate degradation stimulate the rear release of motile vascular smooth muscle cells. J. Biol. Chem. 275, 35384–35392. (2000).

54. W. S. Kamoun et al., Simultaneous measurement of RBC velocity, flux, hematocrit and shear rate in vascular networks. Nat Methods 7, 655–660 (2010).

55. C. M. Leung et al., A guide to the organ-on-a-chip. Nature Reviews Methods Primers 2, 33 (2022).

56. D. Lorusso et al., Practical fabrication of microfluidic platforms for live-cell microscopy. Biomedical microdevices 18, 78 (2016).

