## Supporting Information for "Intussusceptive angiogenesis-on-a-chip: Evidence for transluminal vascular bridging by endothelial delamination"

### Supporting Information Appendix

**Table S1** Peer-reviewed articles containing images of microvessels with transluminal endothelial bridges. Each bridge was classified as: type 1, a thin transluminal filament-like bridge without a nucleus; type 2, a bridge containing part or all of a nucleus; or type 3, a bridge comprised of multiple endothelial cells.

| Number | PMID | First Author (Year) | Context | Number of endothelial bridges displayed | Figure and bridge type displayed |
|---|---|---|---|---|---|
| 1 | 34826235 | Arpino (2021) | Mouse ischemic hindlimb | 7 | Figure 3G: 1<br>Figure 3H: 1, 1<br>Figure 3I: 2, 1<br>Figure 5C: 1<br>Figure 5F: 2 |
| 2 | 12631352 | Notoya (2003) | Rat nephritis | 1 | Figure 6B-C: 1 |
| 3 | 36835203 | Díaz-Flores (2023) | Human Kaposi sarcoma | 29 | Figure 1B: 2, 2, 1, 2, 2, 1<br>Figure 5A: 2, 1<br>Figure 7A: 1, 1, 1<br>Figure 10E: 1, 1, 1, 1, 2, 2, 2<br>Figure 11B: 1, 1<br>Figure 11C: 1, 1, 1<br>Figure 14B: 1, 1, 1, 1, 1<br>Figure 15D: 1 |
| 4 | 34400131 | Pandita (2021) | Human melanoma mouse xenograft | 5 | Figure 2E: 2<br>Figure 3D: 3<br>Figure 4C: 3 |

| | | | | | |
|---|---|---|---|---|---|
| | | | | | Figure 6A-B: 3,2 |
| 5 | 21827961 | Paku (2011) | Mouse colorectal carcinoma line | 4 | Figure 2F: 1<br>Figure 2G: 1, 1, 1 (Net) |
| 6 | 19882213 | Nico (2010) | Human glioblastoma | 6 | Figure 1B: 1, 1, 1<br>Figure 3B: 1, 1, 1 |
| 7 | 20225210 | Konerding (2010) | Murine colitis | 1 | Figure 6A: 1 |
| 8 | 21435466 | Wnuk (2011) | Rat nephritis | 5 | Figure 3E-F: 1, 1, 1, 1, 1 |
| 9 | 35832452 | Kovacs (2022) | Human pleural mesothelioma | 1 | Figure 2D: 1 |
| 10 | 34013885 | Li (2021) | Zebrafish CCM2 knockout | 2 | Figure 3A-B: 3, 1 (Net) |
| 11 | 8654896 | Wilting (1996) | Chick CAM | 3 | Figure 2C: 1, 1<br>Figure 2E: 1 |
| 12 | 11597997 | Patan (2001) | Human tumor xenograft | 2 | Figure 1G: 1<br>Figure 1I: 1 |
| 13 | 15566432 | Fujimoto (2004) | Mouse HT1080 tumour | 1 | Figure 3B: 1 |
| 14 | 19325026 | Makanya (2009) | Embryonic chicken lung | 4 | Figure 5C: 2, 1<br>Figure 5D: 1, 1 |
| 15 | 19555425 | Polykandriotis (2009) | Porous matrix in rats | 1 | Figure 3C: 1 |
| 16 | 31242036 | Ali (2019) | Zebrafish (hypoxic) | 2 | Figure 4C: 1<br>Figure 6F: 1 |
| 17 | 21655651 | Ceauşu (2011) | Liver metastasis from colon carcinoma | 3 | Figure 2D: 1, 1, 1 |
| 18 | 9662652 | Zhou (1998) | Rats treated with prazosin (increased blood flow) | 3 | Figure 4A: 1<br>Figure 4B: 1<br>Figure 4D: 1 |
| 19 | 34930527 | Reichardt (2021) | Human cardiac tissue (COVID-19) | 1 | Figure 4F: 1 |

| | | | | | |
|---|---|---|---|---|---|
| 20 | 36371548 | Werlein (2023) | Human cardiac tissue (COVID-19) | 2 | Figure 6E: 1, 1 |
| 21 | 26310512 | Giacomini (2015) | Murine model of Krabbe disease (frontal cortex) | 2 | Figure 5C: 2, 1 |
| 22 | 27956462 | Xuesong (2016) | Rat glioma | 3 | Figure 2D: 3, 1, 1 |
| 23 | 8781219 | Hansen-Smith (1996) | Rat skeletal muscle (electrical stimulation) | 1 | Figure 4: 1 |
| 24 | 24144103 | Jones (2014) | Adult rat lung subjected to high oxygen | 6 | Figure 1: 2<br>Figure 2A: 2, 2<br>Figure 2B: 2<br>Figure 2F-H: 2, 2 |
| 25 | 7694800 | Wilting (1993) | Chick CAM subjected to VEGF | 6 | Figure 3B: 2, 2, 2<br>Figure 3D: 2, 1<br>Figure 4C: 1 |
| 26 | 12831058 | vanAmerongen (2002) | Subcutaneous foreign body reaction in mice | 2 | Figure 4B: 1, 1 |
| 27 | 29951652 | Zhan (2018) | Implanted porous biomaterials in mice | 5 | Figure 2I: 1<br>Figure 3B: 1<br>Figure 7B-D: 1, 1, 1 |
| 28 | 24011796 | Kılıçaslan (2013) | Cutaneous EGF-exposed wounds in rats | 1 | Figure 8A-B: 1 |
| 29 | 20340003 | Kitahara (2010) | Mouse melanoma | 3 | Figure 8D: 2<br>Figure 8E: 2, 1 (Net) |
| 30 | 30212835 | Logothetidou (2018) | Porcine fetal glomerulus | 5 | Figure 1C-F:1, 1<br>Figure 3A-F: 1, 1<br>Figure 4A-F: 2 |
| 31 | 16003781 | Makanya (2005) | Developing chick kidney | 2 | Figure 4c1-4: 1<br>Figure 4f1-3: 1 |
| 32 | 9056474 | Patan (1997) | Chick CAM | 1 | Figure 4A-D: 1 |

| | | | | | |
|---|---|---|---|---|---|
| 33 | 8238959 | Patan (1993) | Chick CAM | 2 | Figure 4A-D: 1<br>Figure 5A-D: 1 |
| 34 | 35775452 | Díaz-Flores (2023) | Human angiolipoma | 8 | Figure 6A-F: 1, 1, 1, 1, 1, 1, 1, 1 |
| 35 | 34274851 | Taguchi (2021) | Eribulin-induced vascular remodeling in sarcoma xenograft | 1 | Figure 4D: 1 |
| 36 | 33336356 | Díaz-Flores (2021) | Intravascular papillary endothelial hyperplasia | 13 | Figure 2B-D: 1, 1, 1, 1, 1, 1<br>Figure 3A-H:<br>1, 1, 1, 1, 1, 1, 2 |
| 37 | 12203731 | Djonov (2002) | Chick CAM | 1 | Figure 5C-H: 3 |
| 38 | 12926534 | Walski (2003) | Rat cerebral cortex after surgical injury | 2 | Figure 1A: 2<br>Figure 3: 1 |
| 39 | 29643120 | Groppa (2018) | Mouse hindlimb implanted with $VEGFA_{164}$-expressing myoblasts | 11 | Figure 1B: 1<br>Figure 3: 1, 1, 1, 1, 1, 1<br>Figure 4: 1, 1, 2, 1 |
| 40 | 33126763 | Díaz-Flores (2020) | Human gallbladders with cholecystitis | 7 | Figure 1I: 1<br>Figure 2D: 1, 1, 1 (Ner)<br>Figure 4G: 1, 1, 1 (Net) |
| 41 | 34685606 | Díaz-Flores (2021) | Human glioblastoma | 2 | Figure 6C: 1<br>Figure 7I: 1 |
| 42 | 32193418 | Díaz-Flores (2020) | Human lobular capillary hemangioma | 3 | Figure 3M: 1<br>Figure 5E: 1<br>Figure 7L: 2 |
| 43 | 12500160 | Frontczak-Baniewicz (2002) | Traumatic brain injury in rats | 3 | Figure 2: 1<br>Figure 5: 1 |

| | | | | | |
|---|---|---|---|---|---|
| | | | | | Figure 6: 1 |
| 44 | 15944771 | Ribatti (2005) | Human melanoma | 3 | Figure 3A: 1<br>Figure 3B: 1, 1 |
| 45 | 11166277 | Egginton (2001) | Adult rats treated with prazosin | 2 | Figure 4: 1<br>Figure 6: 1 |

CCM2, cerebral cavernous malformation 2; Net, bridge network within a given microvessel; CAM, chorioallantoic membrane; VEGF vascular endothelial growth factor, EGF, epidermal growth factor

**Supporting References for Table S1**


1. Z. Ali et al., Intussusceptive Vascular Remodeling Precedes Pathological Neovascularization. Arterioscler Thromb Vasc Biol 39, 1402-1418 (2019).
2. J.-M. Arpino et al., Low-flow intussusception and metastable VEGFR2 signaling launch angiogenesis in ischemic muscle. Sci Adv 7, eabg9509 (2021).
3. R. A. Ceauşu et al., CD105/Ki67 double immunostaining expression in liver metastasis from colon carcinoma. Rom J Morphol Embryol 52, 613-616 (2011).
4. L. Díaz-Flores et al., Intussusceptive Angiogenesis and Peg-Socket Junctions between Endothelial Cells and Smooth Muscle Cells in Early Arterial Intimal Thickening. Int J Mol Sci 21, 8049 (2020).
5. L. Díaz-Flores et al., Disproportion in Pericyte/Endothelial Cell Proliferation and Mechanisms of Intussusceptive Angiogenesis Participate in Bizarre Vessel Formation in Glioblastoma. Cells 10, 2625 (2021).
6. L. Díaz-Flores et al., Participation of Intussusceptive Angiogenesis in the Morphogenesis of Lobular Capillary Hemangioma. Sci Rep 10, 4987 (2020).
7. L. Díaz-Flores et al., Delimiting CD34+ Stromal Cells/Telocytes Are Resident Mesenchymal Cells That Participate in Neovessel Formation in Skin Kaposi Sarcoma. Int J Mol Sci 24, 3793 (2023).
8. L. Díaz-Flores et al., Myriad pillars formed by intussusceptive angiogenesis as the basis of intravascular papillary endothelial hyperplasia (IPEH). IPEH is intussusceptive angiogenesis made a lesion. Histol Histopathol 36, 217-228 (2021).
9. L. Díaz-Flores et al., Intussusceptive angiogenesis facilitated by microthrombosis has an important example in angiolipoma. An ultrastructural and immunohistochemical study. Histol Histopathol 38, 29-46 (2023).

10. V. G. Djonov, H. Kurz, P. H. Burri, Optimality in the developing vascular system: branching remodeling by means of intussusception as an efficient adaptation mechanism. Dev Dyn 224, 391-402 (2002).

11. S. Egginton, A. L. Zhou, M. D. Brown, O. Hudlická, Unorthodox angiogenesis in skeletal muscle. Cardiovasc Res 49, 634-646 (2001).

12. M. Frontczak-Baniewicz, M. Walski, Non-sprouting angiogenesis in neurohypophysis after traumatic injury of the cerebral cortex. Electron-microscopic studies. Neuro Endocrinol Lett 23, 396-404 (2002).

13. A. Fujimoto et al., Vascular endothelial growth factor reduces mural cell coverage of endothelial cells and induces sprouting rather than luminal division in an HT1080 tumour angiogenesis model. Int J Exp Pathol 85, 355-364 (2004).

14. A. Giacomini et al., Brain angioarchitecture and intussusceptive microvascular growth in a murine model of Krabbe disease. Angiogenesis 18, 499-510 (2015).

15. E. Groppa et al., EphrinB2/EphB4 signaling regulates non-sprouting angiogenesis by VEGF. EMBO Rep 19, e45054 (2018).

16. F. M. Hansen-Smith, O. Hudlicka, S. Egginton, In vivo angiogenesis in adult rat skeletal muscle: early changes in capillary network architecture and ultrastructure. Cell Tissue Res 286, 123-136 (1996).

17. R. C. Jones, D. E. Capen, Mechanisms of growth of a pulmonary capillary network in adult lung. Ultrastruct Pathol 38, 34-44 (2014).

18. Z. K et al., Different angiogenesis modes and endothelial responses in implanted porous biomaterials. Integrative biology : quantitative biosciences from nano to macro 10 (2018).

19. S. Kitahara, S. Morikawa, K. Shimizu, H. Abe, T. Ezaki, Alteration of angiogenic patterns on B16BL6 melanoma development promoted in Matrigel. Med Mol Morphol 43, 26-36 (2010).

20. S. M. S. Kılıçaslan, S. C. Cevher, E. G. G. Peker, Ultrastructural changes in blood vessels in epidermal growth factor treated experimental cutaneous wound model. Pathol Res Pract 209, 710-715 (2013).

21. M. A. Konerding et al., Inflammation-induced intussusceptive angiogenesis in murine colitis. Anat Rec (Hoboken) 293, 849-857 (2010).

22. I. Kovacs et al., Malignant pleural mesothelioma nodules remodel their surroundings to vascularize and grow. Transl Lung Cancer Res 11, 991-1008 (2022).

23. W. Li et al., Abortive intussusceptive angiogenesis causes multi-cavernous vascular malformations. Elife 10, e62155 (2021).

24. A. Logothetidou et al., Intussusceptive Pillar Formation in Developing Porcine Glomeruli. J Vasc Res 55, 278-286 (2018).

25. A. N. Makanya, V. Djonov, Parabronchial angioarchitecture in developing and adult chickens. J Appl Physiol (1985) 106, 1959-1969 (2009).

26. A. N. Makanya, D. Stauffer, D. Ribatti, P. H. Burri, V. Djonov, Microvascular growth, development, and remodeling in the embryonic avian kidney: the interplay between sprouting and intussusceptive angiogenic mechanisms. Microsc Res Tech 66, 275-288 (2005).

27. B. Nico et al., Intussusceptive microvascular growth in human glioma. Clin Exp Med 10, 93-98 (2010).

28. M. Notoya, T. Shinosaki, T. Kobayashi, T. Sakai, H. Kurihara, Intussusceptive capillary growth is required for glomerular repair in rat Thy-1.1 nephritis. Kidney Int 63, 1365-1373 (2003).

29. S. Paku et al., A new mechanism for pillar formation during tumor-induced intussusceptive angiogenesis: inverse sprouting. Am J Pathol 179, 1573-1585 (2011).

30. A. Pandita et al., Intussusceptive Angiogenesis in Human Metastatic Malignant Melanoma. Am J Pathol 191, 2023-2038 (2021).

31. S. Patan, B. Haenni, P. H. Burri, Evidence for intussusceptive capillary growth in the chicken chorio-allantoic membrane (CAM). Anat Embryol (Berl) 187, 121-130 (1993).

32. S. Patan, B. Haenni, P. H. Burri, Implementation of intussusceptive microvascular growth in the chicken chorioallantoic membrane (CAM). Microvasc Res 53, 33-52 (1997).

33. S. Patan et al., Vascular morphogenesis and remodeling in a human tumor xenograft: blood vessel formation and growth after ovariectomy and tumor implantation. Circ Res 89, 732-739 (2001).

34. E. Polykandriotis et al., Regression and persistence: remodelling in a tissue engineered axial vascular assembly. J Cell Mol Med 13, 4166-4175 (2009).

35. M. Reichardt et al., 3D virtual histopathology of cardiac tissue from Covid-19 patients based on phase-contrast X-ray tomography. Elife 10, e71359 (2021).

36. D. Ribatti et al., Microvascular density, vascular endothelial growth factor immunoreactivity in tumor cells, vessel diameter and intussusceptive microvascular growth in primary melanoma. Oncol Rep 14, 81-84 (2005).

37. E. Taguchi et al., Eribulin induces tumor vascular remodeling through intussusceptive angiogenesis in a sarcoma xenograft model. Biochem Biophys Res Commun 570, 89-95 (2021).

38. M. J. van Amerongen, G. Molema, J. Plantinga, H. Moorlag, M. J. A. van Luyn, Neovascularization and vascular markers in a foreign body reaction to subcutaneously implanted degradable biomaterial in mice. Angiogenesis 5, 173-180 (2002).

39. M. Walski, M. Frontczak-Baniewicz, New vessel formation after surgical brain injury in the rat's cerebral cortex II. Formation of the blood vessels distal to the surgical injury. Acta Neurobiol Exp (Wars) 63, 77-82 (2003).

40. C. Werlein et al., Inflammation and vascular remodeling in COVID-19 hearts. Angiogenesis 26, 233-248 (2023).

41. J. Wilting et al., VEGF121 induces proliferation of vascular endothelial cells and expression of flk-1 without affecting lymphatic vessels of chorioallantoic membrane. Dev Biol 176, 76-85 (1996).

42. J. Wilting, B. Christ, M. Bokeloh, H. A. Weich, In vivo effects of vascular endothelial growth factor on the chicken chorioallantoic membrane. Cell Tissue Res 274, 163-172 (1993).

43. M. Wnuk, R. Hlushchuk, G. Tuffin, U. Huynh-Do, V. Djonov, The effects of PTK787/ZK222584, an inhibitor of VEGFR and PDGFRβ pathways, on intussusceptive angiogenesis and glomerular recovery from Thy1.1 nephritis. Am J Pathol 178, 1899-1912 (2011).

44. D. Xuesong et al., Evaluation of neovascularization patterns in an orthotopic rat glioma model with dynamic contrast-enhanced MRI. Acta Radiol 58, 1138-1146 (2017).

45. A. Zhou, S. Egginton, O. Hudlická, M. D. Brown, Internal division of capillaries in rat skeletal muscle in response to chronic vasodilator treatment with alpha1-antagonist prazosin. Cell Tissue Res 293, 293-303 (1998).

**Table S2** Demographics of patients with skin wounds due to peripheral artery disease necessitating lower limb amputation

| Patient | Age | Sex | Diabetes | Ulcer site | Surgical procedure |
|---|---|---|---|---|---|
| 1 | 46 | F | yes | Right great toe and forefoot | Below-knee amputation |
| 2 | 56 | M | yes | Right forefoot | Below-knee amputation |
| 3 | 60 | M | yes | Right stump wound | Above-knee amputation* |
| 4 | 70 | M | yes | Left great toe | Below-knee amputation |
| 5 | 73 | M | yes | Right heel | Below-knee amputation |
| 6 | 74 | M | no | Right heel | Below-knee amputation |
| 7 | 79 | M | yes | Left great toe | Below-knee amputation |
| 8 | 86 | M | yes | Left forefoot | Below-knee amputation |
| 9 | 88 | M | yes | Right foot | Above-knee amputation |

*Subsequent to a below-knee amputation

A

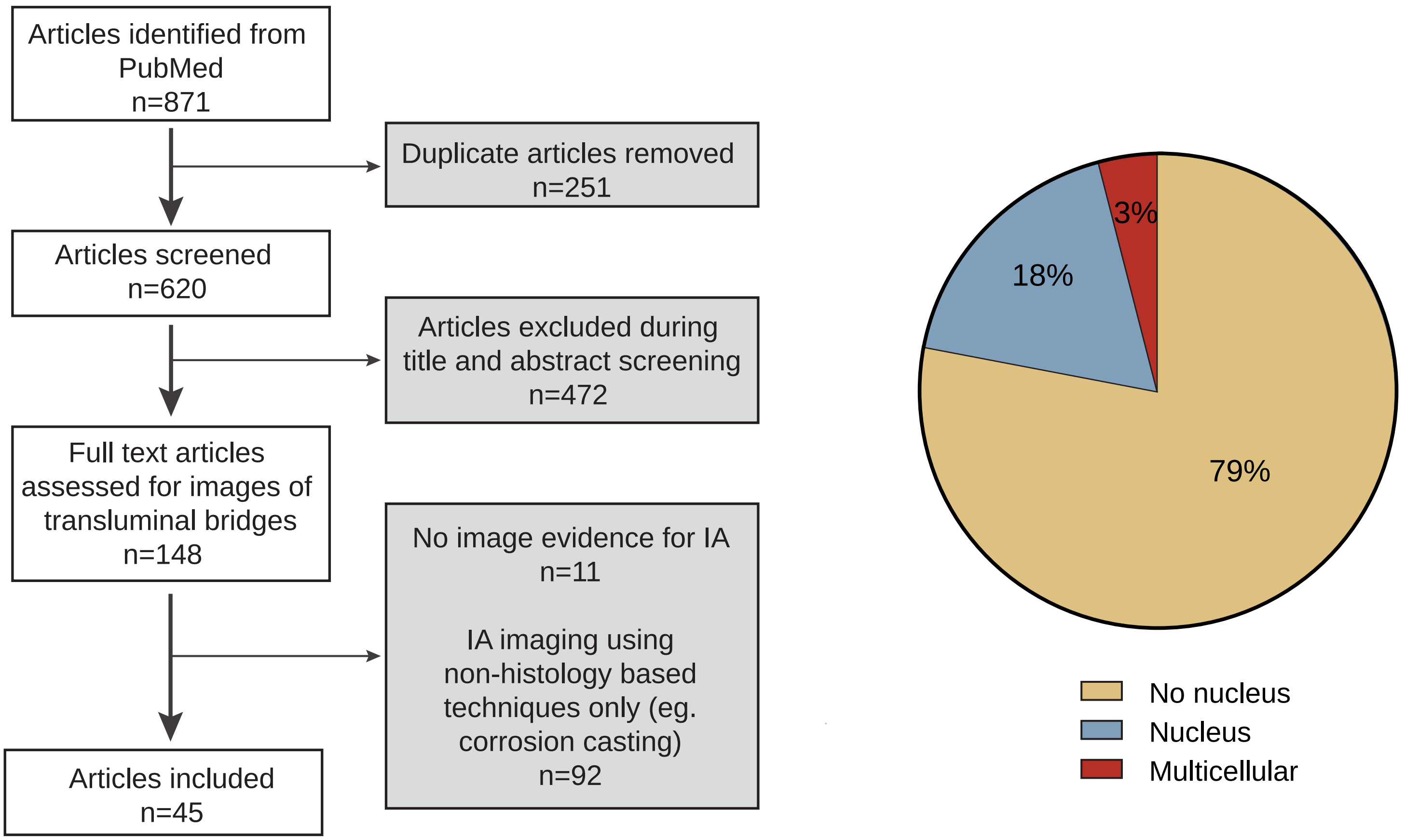


B

| Bridge type | Article count | Image count | Bridge count | Context examples |
|---|---|---|---|---|
| No nucleus (Type 1) | 42 | 94 | 141 | Chick CAM (Patan et al., 1997)<br>Rat nephritis (Notoya et al., 2003)<br>Rat cerebral cortex after surgical injury (Walski et al., 2003)<br>Porcine fetal glomerulus (Logothetidou et al., 2018)<br>Human pleural mesothelioma (Kovacs et al., 2022) |
| Nucleus (Type 2) | 13 | 21 | 31 | Embryonic chicken lung (Makanya et al., 2009)<br>Murine model of Krabbe disease (Giacomini et al., 2015)<br>Adult rat lung after injury in high oxygen (Jones et al., 2014) |
| Multicellular (Type 3) | 4 | 6 | 6 | CCM2 KO vascular lesion in zebrafish (Li et al., 2021)<br>Rat glioma (Xuesong et al., 2017)<br>Human melanoma xenograft (Pandita et al., 2021) |
| Network | 5 | 5 | 5 | Murine experimental tumors (Paku et al., 2011)<br>Human angiolipoma (Díaz-Flores et al., 2023)<br>Human gallbladders with acute cholecystitis (Díaz-Flores et al., 2020) |

**Figure S1. Systematic review of the morphology of transluminal endothelial bridges**

**A.** Protocol for systematically identifying and assessing published images of transluminal endothelial bridges. Only endothelial cell bridges were evaluated, not multi-component pillars or flow dividers of later-stage IA. **B.** Table showing examples of the endothelial bridge categories arising from the analysis is shown. These categories are homologous to those found in the micro-physiologic system. References can be found in Supporting References for Table S1.

**Legends For Movies**

**Movie S1:** Three-dimensional volume projections of endothelial cells lining a fibronectin-coated PDMS channel subjected to 2 dyn/cm$^2$ shear stress stained with Alexa Fluor-conjugated phalloidin. A cord-like transluminal bridge is seen.

**Movie S2**: Three-dimensional volume projections of endothelial cells lining a fibronectin-coated PDMS channel subjected to 2 dyn/cm$^2$ shear stress stained with Alexa Fluor-conjugated phalloidin. A cord-like transluminal bridge is seen with a cell nucleus within the bridge, seemingly entering from the wall-bridge junction.

**Movie S3:** Three-dimensional volume projections of endothelial cells in the microfluidic channel subjected to 2 dyn/cm$^2$ stained with Alexa Fluor-conjugated phalloidin. A multicellular cylindrical bridge can be seen to transect the lumen. The bridge itself has bifurcations.

**Movie S4:** Time-lapse video sequence of GFP-labeled endothelial cells lining a microfluidic device. The lower component of the channel is shown, and cells are color-mapped based on their vertical distance from the bottom of the channel. A circumferentially aligned endothelial cell, i.e. aligned orthogonal to the direction of flow, is denoted by the arrow. This cell protrudes and part of the stretched tail delaminates to form a bridge (arrow).

**Movie S5:** Time-lapse video sequence of a microvessel-on-a-chip showing an endothelial cell crawling along the side wall of the channel (arrowhead), sending a protrusion orthogonal to the direction of flow (arrow), which then delaminates to form a bridge (arrow). The movie is briefly paused when arrows appear, to facility tracking the motility events.

**Movie S6:** Time-lapse video sequence of a microvessel-on-a-chip with endothelial cells stained with LIVE 610-conjugated jasplakinolide. Two adjacent endothelial cells (arrows) can be seen circumferentially crawling in opposite directions while remaining attached to each other. The stretch cell-cell connection delaminates off the bottom surface of the channel to form a bridge (arrow).

**Movie S7:** Time-lapse video sequence of a microvessel-on-a-chip lined with GFP-expressing endothelial cells. A circumferentially oriented endothelial cell can be seen to release from an adjacent, similarly aligned cell. The cell ends remain attached to the bottom and lateral surfaces of the channel, and the intervening segment snaps into a bridging position (arrow).

**Movie S8:** Time-lapse video sequence of a microvessel-on-a-chip lined with GFP-expressing endothelial cells with a transluminal bridging extending from a sidewall to the bottom surface. The bridge connection at the bottom is dynamically remodeled so that the bridge effectively walks from one endothelial cell to an adjacent one.
